# Diversity of mycotoxins and other secondary metabolites recovered from blood oranges infected by *Colletotrichum*, *Alternaria*, and *Penicillium* species

**DOI:** 10.1101/2023.05.09.540008

**Authors:** Ermes Ivan Rovetto, Carlos Luz, Federico La Spada, Giuseppe Meca, Mario Riolo, Santa Olga Cacciola

## Abstract

This study identified secondary metabolites produced by *Alternaria alternata*, *Colletotrichum gloeosporioides* and *Penicillium digitatum* in fruits of two blood orange cultivars before harvest. Analysis was performed by UHPLC–Q-TOF-MS. Three types of fruits were selected, asymptomatic, symptomatic showing necrotic lesions caused by hail, and mummified. Extracts from peel and juice were analyzed separately. *Penicillium digitatum* was the prevalent species recovered from mummified and hail-injured fruits. Among 47 secondary metabolites identified, 16, 18 and 13 were of *A. alternata*, *C. gloeosporioides* and *P. digitatum*, respectively. Consistently with isolations, indicating the presence of these fungi also in asymptomatic fruits, the metabolic profiles of the peel of hail-injured and asymptomatic fruits did not differ substantially. Major differences were found in the profiles of juice from hail injured and mummified fruits, such as a significant higher presence of 5,4-dihydroxy-3,7,8-trimethoxy-6C-methylflavone and Atrovenetins, particularly in the juice of mummified fruits of Tarocco Lempso cultivar. Moreover, the mycotoxins Patulin and Rubratoxin B were detected exclusively in mummified fruits. Patulin was detected in both the juice and peel, with a higher relative abundance in the juice, while Rubratoxin B was detected only in the juice. These findings provide basic information for evaluating and preventing the risk of contamination by mycotoxins in the citrus fresh fruit supply chain and juice industry.

## 1. Introduction

The genus *Citrus*, family *Rutaceae*, comprises some of the most widely cultivated fruit crops worldwide, such as oranges, lemons, tangerines, mandarins, limes, grapefruits and citrons [1–3]. Globally, citrus are cultivated in tropical, subtropical and temperate climates in more than 140 countries and are consumed mainly as fresh fruit or juice [4]. Blood oranges are a group of sweet orange (*Citrus* x *sinensis*) cultivars characterized by a red pigmentation of the rind and flesh, of variable intensity. These oranges are appreciated for their organoleptic characteristics and richness in anthocyanins, a family of polyphenolic pigments. The health benefits of anthocyanins, which possess strong antioxidant properties, include antidiabetic, anticancer, anti-inflammatory, antimicrobial and anti-obesity effects as well as the prevention of cardiovascular diseases [5]. Like all citrus fruits, blood oranges are a source of other health-promoting substances, such as vitamins, mineral salts, phenolic acids, flavonoids, pectin and dietary fibers [1,6,7]. The traditional growing areas of blood oranges have long been Sicily, still the leading producer in the world, Spain, Morocco and Tunisia. As a typical local product, in the EU the red orange from Sicily has Protected Geographical Status. Among the Sicilian blood orange types the most popular is ‘Tarocco’ which comprises numerous cultivars, differing in intensity of red pigmentation and ripening season, fruit shape and size, tree vigor and productivity [8]. In the last years, blood oranges have gained vast popularity and their consumption as fresh fruit or juice has increased and is extending to new countries [9].

Citrus fruits in general are susceptible to infections of fungal pathogens, such as species of *Alternaria*, *Colletotrichum*, *Geotrichum* and *Penicillium* [10–13], which are responsible for both pre- and post-harvest rots. The most common and serious post-harvest rots of citrus fruits are green and blue molds caused by *Penicillium digitatum* and *P. italicum*, respectively, followed by sour rot caused by *Geotrichum citri-aurantii* [14,15]. These fungi are strict wound pathogens and can infect the fruit in the grove, in the packinghouse, or during subsequent handling and storage. *Colletotrichum gloeosporioides* is responsible for both pre- and post-harvest citrus fruit anthracnose and is the prevalent *Colletotrichum* species associated to citrus worldwide [11,16–18]. It resides in citrus fruit peel, supposedly as a saprophyte, endophyte or epiphyte [19], and can switch to a pathogenic lifestyle under the effect of abiotic stress factors, such as exposure to ethylene during post-harvest fruit degreening or injuries caused by wind, heat, frost or hail in the field. The precise identification of *Alternaria* species associated with citrus fruits is problematic, as the taxonomy of *Alternaria* is not well defined due to the great morphological and genetic variability of presently recognized species [20–22]. *Alternaria alternata sensu* Woundeberg et al. [20] is the prevalent among the small-spored *Alternaria* species associated with brown spot of citrus fruits. Brown spot is a disease affecting mainly tangerines and their hybrids, and occurring prevalently in citrus production areas with a Mediterranean type climate [16,23-25]. Moreover, it is the prevalent species associated with Alternaria black rot of citrus fruits, also known as core rot [26,27]. Both of these diseases seriously affect the marketability of fresh citrus fruits. Like *C. gloeosporioides*, *A. alternata* and other small-spored *Alternaria* species reside in the citrus fruit peel as latent pathogens and can infect and colonize, as opportunistic necrotrophic pathogens, injured fruits of all citrus varieties, including those resistant to the Alternaria brown spot disease. Fungal pathogens of citrus fruits were reported to produce mycotoxins [3,24,28,29].

The term mycotoxins refers to secondary metabolites toxic for humans and animals produced by filamentous fungi that colonize crop plants as pathogens or saprophytes [28]. Some fungal genera in particular, including *Alternaria*, *Aspergillus*, *Fusarium*, *Claviceps*, and *Penicillium*, are known to produce mycotoxins [29-31,32]. The most studied mycotoxins include: aflatoxins (AFs), common in dried fruits and nuts; ochratoxin A (OTA), occurring in grapes, raisins and red wines; citrinin (CIT), mainly found in stored grains, but occasionally also in fruits and other plant products. Mycotoxins, including zearalenone (ZEA), fumonisins (FBs) and deoxynivalenol (DON), which is a mycotoxin of the group of trichothecenes (TCTs), are produced by species of *Fusarium* and occurring in cereal crops. Moreover, patulin (PAT), a polyketide mycotoxin produced primarily by *P. expansum* in rotten apples and also occurring in apple juices. Furthermore, ergot alkaloids, indole compounds, are produced by various *Claviceps* species on monocotyledonous plants of the families *Poaceae*, *Juncaceae* and *Cyperaceae*, including forage grasses, corn, wheat, barley, oats, millet, sorghum, rice, and rye. Finally, the dibenzopyrone derivative alternariol (AOH) is produced by the genus *Alternaria* and is a common contaminant of many fruits and vegetables [28,29,33–36]. Beside AOH, *Alternaria* mycotoxins (ATs) comprise alternariol monomethyl ether (AME) and altenuene (ALT), both of the structural group of the dibenzopyrone derivatives, the perylene derivatives altertoxins (ATX-I, ATX-II, and ATX II), and the tetramic acid derivative tenuazonic acid (TeA) [31,34,37–40]. Levels of contamination of plant products by mycotoxins may vary depending on several factors, such as cultivar susceptibility, climate, storage condition, damages caused by insect pests, use of pesticides, geographical origin and mechanical injures due to improper handling or weather events [40,41]. Most mycotoxins are heat stable and resist industrial processes, consequently they can be found both in fresh fruits and processed products, such as fruit juices [40]. Various mycotoxins have been identified in apple, apricot, berry, orange, peach and pear juices [31,41–46].

Contamination by mycotoxins of plant products and processed food and feed are a serious concern for human and animal health and several countries have set regulatory limits for mycotoxins in foods and feeds [24,47–49]. Moreover, strategies for minimizing contamination by mycotoxins in foods and feeds have been suggested [30]. Finally, actions have been taken to prevent or reduce the risk posed by mycotoxins for human health. In particular, the European Commission (EC) has reinforced border controls for mycotoxins in specific crops imported from given geographical areas [50–52]. European and National regulations have established maximum tolerable levels of principal mycotoxins in foods [52]. In fruits and juices, only PAT and OTA are regulated [41]. For PAT, the European Union (EU) [53] set a maximum level of 50 µg/kg in fruit juices, concentrated fruit juices, fruit nectars, cider and other fermented drinks derived from apples or containing apple juice [3,54]. A limit of 25 µg/kg has been implemented in processed solid apple products (apple compote, apple puree intended for direct consumption) [38,54] and 10 µg/kg for apple-derived products intended for young children and infants [38,54]. For OTA, the EC has set 2 μg/kg as maximum level in reconstituted concentrated grape juice [41], grape nectar [40], grape must and reconstituted concentrated grape must, intended for direct human consumption [53]. The European Union (EU) has set the maximum acceptable level for Aflatoxin B1 (AFB1) at 2 μg/kg–12 μg/kg (depending on the type of foodstuff) [55]. The EC Recommendation 2006/576/EC5 [56] regulates the presence of DON, ZEA, OTA, FBs, T-2 and HT-2, the last two both of the TCTs group, in products intended for animal feeding [57]. By contrast, even though EFSA (European Food Safety Authority) has published a scientific opinion on the risks deriving for the animal and human health from the presence of ATs in food and feed [38,58], no EU decision has been so far issued to regulate other group of mycotoxins, such as ATs [38,58].

Both the concern and awareness of the risks posed by mycotoxins to the health safety prompted to seek for new robust analytical methods for the extraction and determination of these metabolites in plant products and processed foods [57]. The most common methods for the extraction of mycotoxins in foodstuffs are QuEChERS extraction [41,59], liquid-liquid extraction [41,60], and dispersive liquid-liquid microextraction (DLLME) [40,41,61]. In particular, the dispersive liquid–liquid microextraction (DLLME) has showed some advantages, such as simplicity of operation [61], high recoveries, low-cost applications [40,41,61,62] and automation of the analytical procedure [40,63]. This method has been used for the determination of many mycotoxins, for example OTA in wines [61,64], ZEA in beer [64,65], AFs in oils, PAT in apple juices, AFs and OTA in rice [61]. Tolosa et al. [66] used the DLLME for multimycotoxins analysis to evaluate the presence of 15 mycotoxins in water and fish plasma [64]. Rodríguez-Carrasco et al. [67] utilized this method for the analysis of ATs in tomato and tomato-based products and Serrano and colleagues [68] used it to investigate the enniatins migration from pasta to pasta cooking water.

The most common techniques used for determination of mycotoxins in food and feed are Enzyme-linked immunosorbent assay (ELISA), High-performance liquid chromatography (HPLC), liquid chromatography-tandem mass spectrometry (LC199 MS/MS), gas chromatography (GC), thin-layer chromatography (TLC) [32,37,69,70] and Ultra Performance Liquid Chromatography (UPLC) [71]. Among them, Liquid chromatography (LC) matched with mass spectrometry (MS) offers a rapid, cost-effective and precise detection of mycotoxins [41,70,71].

Some authors reported HPLC technique application for the analysis of mycotoxins in feeds [72–75], fungal cultures [76], cheese [77], milk [78], bee pollen [79], cereal products [72,80,81] beer and wine [82,83]. In recent years, High-performance liquid chromatography quadrupoletime-of-flight mass spectrometry (UHPLC– Q-TOF-MS) has been evidenced to be a smart and efficient analytical method with high sensitivity, high precision, high resolution and fast information acquisition [84]. The HPLC-Q-TOF-MS has been utilized to study changes induced by bio-preservative microrganisms in fungal metabolomic profiles of food contaminant fungi [85], to determine pesticides on tomato peels [86], and to analyze polymethoxylated flavonoids (PMFs) in citrus peels [87] and plants [84].

The aims of this study were: (*i*) to apply the UHPLC– Q-TOF-MS analytical method for the characterization of the mycotoxicological profile of mature fruits of two blood orange cultivars; (*ii*) to determine how this profile varies depending on the fungi associated to the fruit peel and in the diverse parts of the fruit; (*iii*) to investigate how the profile is affected by the phytosanitary status of the fruit as a consequence of an environmental stress, such as hail.

## 2. Results

Approximately 2800 fungal mass isolates were obtained from orange fruits. Mass isolates were preliminarily separated into three groups on the basis of morphotype, i.e. colony morphology on PDA and microscopical traits. The three morphotypes corresponded to *Alternaria*, *Colletotrichum* and *Penicillium,* respectively. On PDA, *Alternaria* isolates showed flat, woolly colonies ranging from dark green to black in color. They produced typical dark brown, club-shaped conidia arranged in branched chains, with oval-ellipsoidal shapes and 3–5 transverse septa. Colonies of *Colletotrichum* isolates grown on PDA showed a dense, cottony, aerial mycelium that was initially white and turned progressively pale grey with salmon-pink conidial mucilaginous masses in the center of the colony. Dark acervuli were scattered over the entire surface of old colonies, and the colony reverse was pale orange to uniformly grey. The single-celled conidia were hyaline, smooth, cylindrical, with both ends rounded, and dimensions of 11–15 × 4–6 μm. Setae were present in most isolates. On PDA, colonies of *Penicillium* isolates were initially white, and then turned green due to abundant production of conidia. The colony margin was entire and narrow, and conidia were borne in chain on typically branched conidiophores; they were globose to subglobose in shape, smooth, with a size range of 2.51–4.22 × 2.35–3.61 μm and mean size (± SD) of 3.38 ± 0.49 × 3 ± 0.36 μm.

Isolates of *Alternaria*, *Colletotrichum* and *Penicillium*, accounted for 12, 28 and 60% of all isolates recovered from orange fruits, respectively, and the proportions of isolates of the three fungal genera in each of the six clusters of fruits, as defined in Table 3, are reported in Table 1.

**Table 1.**
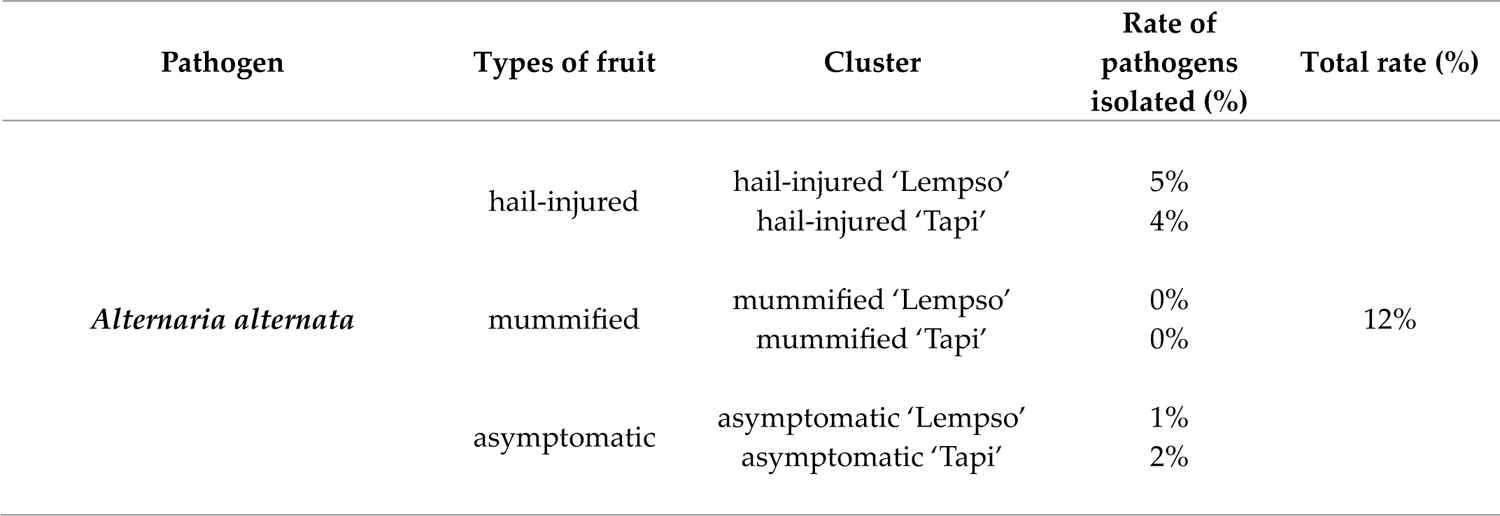
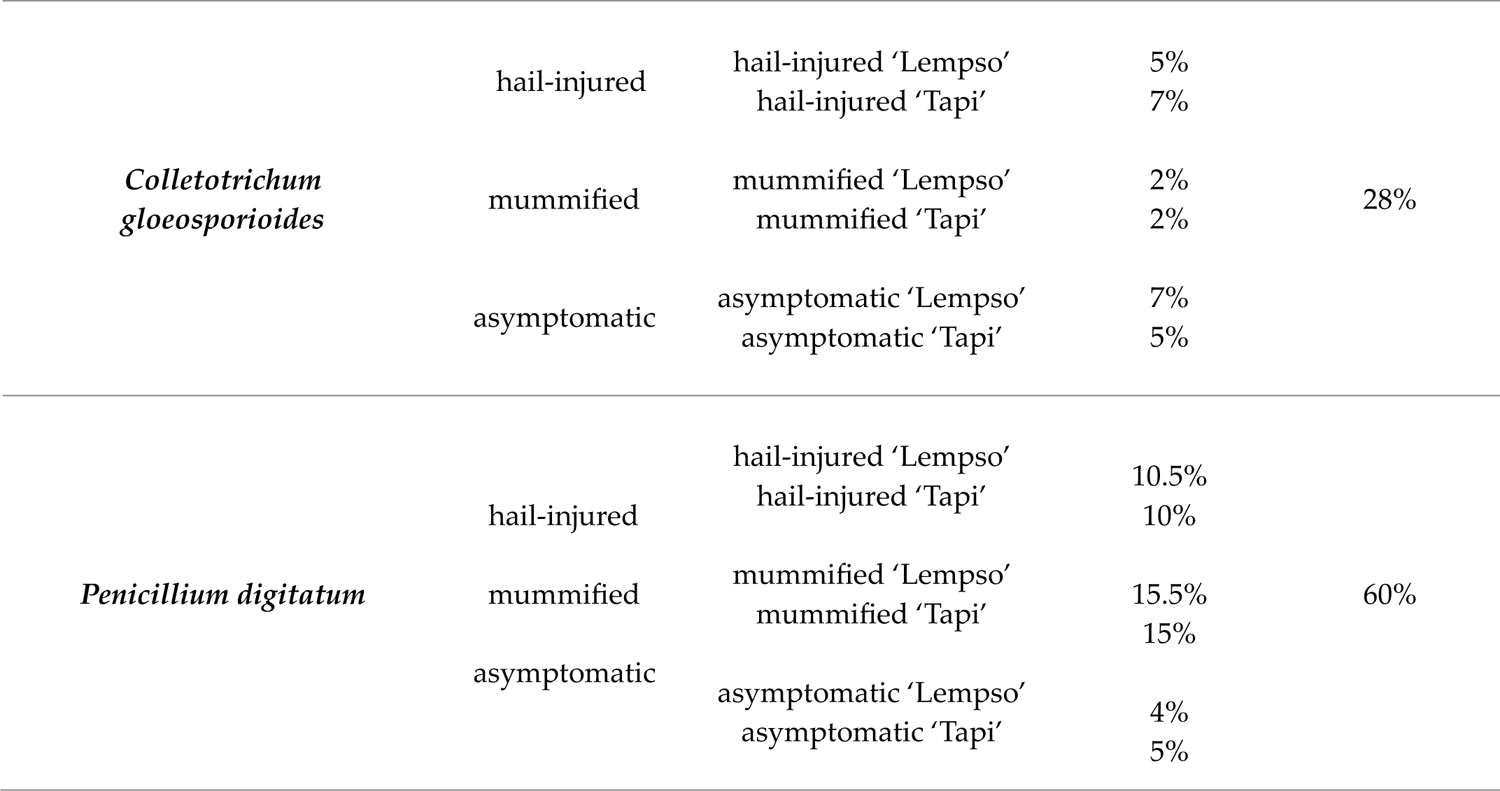
Proportion of isolates of three fungal species from three different types of orange fruits subdivided into proportion of isolates per each cluster. A set of about 7% of mass isolates of each fungal genus, comprising isolates from all clusters, was randomly selected and a single-conidium isolate was obtained from each selected mass isolate to be identified molecularly. Overall, 200 isolates, 24 of *Alternaria*, 56 of *Colletotrichum* and 120 of *Penicillium*, were identified molecularly at species level. In particular, based on the phylogenetic analysis of both the ITS and βtub regions the *Colletotrichum* and *Penicillium* isolates were identified as *C. gloeosporioides* and *P. digitatum*, respectively, while based on the phylogenetic analysis of ITS, TEF-1a and Alt-1a regions the *Alternaria* isolates were identified as *A. alternata*. As the isolates belonging to the same species resulted to be identical, a limted number of sequences of isolates per each species were deposited on Genbank (Table 2).

The analysis by Agilent Ultra High-Definition Accurate Mass revealed 47 diverse fungal secondary metabolites in peel and juice of orange fruits. The identification of the metabolites was supported by using a specifically designed database based on the secondary metabolites produced by species of *Alternaria*, *Colletotrichum* and *Penicillium* available in the literature (Table 3).

The secondary metabolites detected by UHPLC– Q-TOF-MS in each fruit cluster are presented as heat maps (Figures 1 - 3). Only secondary metabolites with a large gap between the asymptomatic fruit cluster of each orange cultivar and the clusters of both hail-injured and mummified fruits of the same cultivar are reported in the maps.

**Figure 1.**
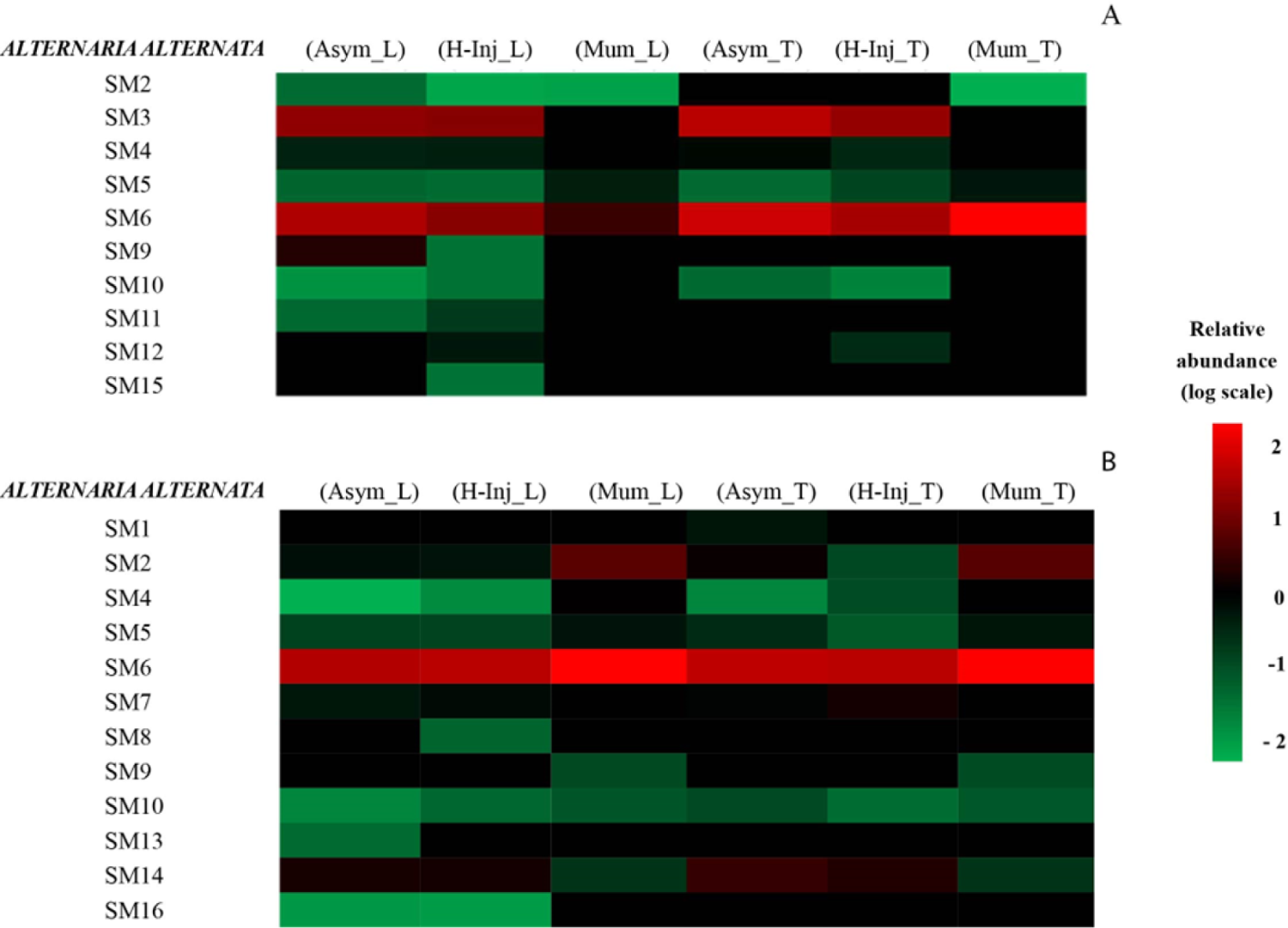
Heat map representing the relative abundance of *Alternaria alternata* secondary metabolites detected in orange peel (A) and orange juices (B). Colors are based on the relative abundance (log-arithmic scale) of the secondary metabolites detected, where red represents high abundance and green represents low abundance.

Figure 1A shows that the secondary metabolites of *A. alternata* present in orange peel included SM2, SM3, SM4, SM5, SM6, SM9, SM10, SM11, SM12 and SM15. However, only SM3 (AF-toxins II) and SM6 (Alternethanoxin A) were found with a high relative abundance compared to the other metabolites. Specifically, SM3 was found with a high relative abundance in all clusters except for the mummified fruits of both varieties, while SM6 was relatively abundant in all clusters, except in Mum_L.

SM2 (AAL-toxin TE2) was present only in (Mum_T) cluster; in (Mum_L) cluster, SM5 (Alternariol monomethyl ether, AME) had a relative abundance, approximately 6-fold higher than the (Asym_L) cluster. In addition, in (H-Inj_T) and (Mum_T) clusters, this metabolite had relative abundance approximately 3-fold and 9-fold higher than (Asym_T) cluster. Alternariol monomethyl ether (AME) (SM5) was report as cytotoxic activity [90]. In (Mum_T) cluster, SM6 had relative abundance approximately 2-fold higher than in (Asym_T) cluster. Alternethanoxin A (SM6) displayed *in vitro* growth inhibitory activity with respect to several cancer cell lines [123]. Furthermore, SM10 (Dihydroaltersolanol) and SM11 (Erythroglaucin), in (H-Inj_L) had relative abundance approximately 2-fold and 3-fold higher than in (Asym_L).

Figure 1B shows that secondary metabolites released by *A. alternata* in orange juice were SM1, SM2, SM4, SM5, SM6, SM7, SM8, SM9, SM10, SM13, SM14 and SM16.

However, only SM2, SM6 and SM14 (Porriolide) were found to be relatively abundant compared to the other metabolites. Specifically, in mummified fruits of both orange cultivars, (Mum_L) and (Mum_T) clusters, SM2 showed a relative abundance around 4- and 3-fold higher, respectively, than in asymptomatic fruits, (Asym_L) and (Asym_T) clusters. In hail-injured fruits of both orange cultivars, (H-Inj_L) and (H-Inj_T) clusters, SM4 (Altenusin) showed a relative abundance around 3- and 6-fold higher than in asymptomatic fruits, (Asym_L) and (Asym_T) clusters, respectively. In mummified fruits of both orange cultivars, (Mum_L) and (Mum_T) clusters, this metabolite showed a relative abundance about 251- and 63-fold higher than in asymptomatic fruits, (Asym_L) and (Asym_T) clusters, respectively. In (Mum_L) and (Mum_T) clusters, SM5 showed a relative abundance about 4- and 2-fold higher than in (Asym_L) and (Asym_T) clusters, respectively. In (Mum_L) and (Mum_T) clusters, SM6 (Alternethanoxin A) showed a relative abundance about 2-fold higher than in (Asym_L) and (Asym_T) clusters, respectively. SM9 (Curvularin) was present only in (Mum_L) and (Mum_T) clusters, while SM10 (Dihydroaltersolanol) in (H-Inj_L) and (Mum_L) clusters showed a relative abundance approximately 2- and 5-fold higher, respectively, than in (Asym_L). SM14 (Porriolide) was recorded with a high relative abundance in (Asym_T) and (H-Inj_T) compared with other clusters.

Figures 2A and B show the relative abundance of secondary metabolites of *Colletotrichum* produced in the peel and juice of orange fruits. The following metabolites were detected in the peel: SM19, SM20, SM21, SM22, SM24, SM25, SM27, SM28, SM29, SM30, SM31, SM32, and SM33. Among them, SM19 (5,4-dihydroxy-3,6,7-trimethoxy-8C-methylflavone) showed a higher relative abundance in the hail-injured fruits, (H-Inj_L) cluster, compared to the asymptomatic fruits, (Asym_L) cluster. Similarly, the (H-Inj_T) and (Mum_T) clusters showed a higher relative abundance in SM19 (5,4-dihydroxy-3,6,7-trimethoxy-8C-methylflavone) than in the (Asym_T) cluster. SM20 (5,4-dihydroxy-3,7,8-trimethoxy-6C-methylflavone) and SM22 (Apigenin-8-C-β-D-glucopyranoside) were found with a high relative abundance in all clusters of both varieties. SM21 (Alternariol-5-Me ether) showed a notable higher relative abundance in the (Mum_L) cluster compared to the (Asym_L) cluster. SM21 (Alternariol-5-Me ether) showed a higher relative abundance in the (H-Inj_T) and (Mum_T) clusters compared to the (Asym_T) cluster. SM25 (Colletoic acid) was detected at higher levels in both (Mum_L) and (Mum_T) clusters compared to the (Asym_L) and (Asym_T) clusters. In (Mum_L) and (Mum_T) clusters, SM29 (Colletotrilactam A) was detected with a higher relative abundance than in other clusters.

**Figure 2.**
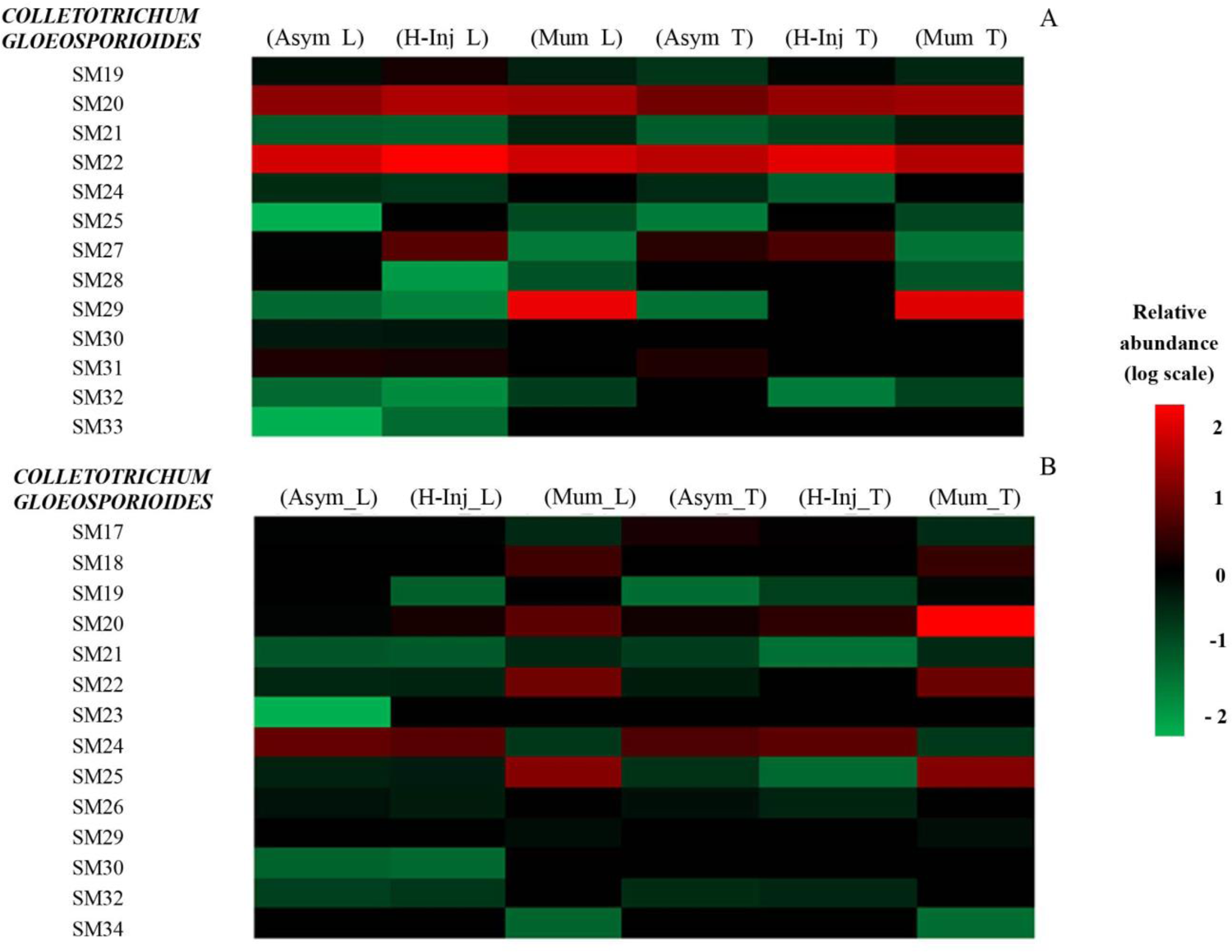
Heat map representing the relative abundance of *Colletotrichum gloeosporioides* secondary metabolites detected in orange peel (A) and orange juices (B). Colors represent the relative abundance (logarithmic scale) of the secondary metabolites: red represents high abundance, green low abundance.

Figure 2B shows the relative abundance of secondary metabolites produced by *C. gloeosporioides* in orange juice, including SM17, SM18, SM19, SM20, SM21, SM22, SM23, SM24, SM25, SM26, SM29, SM30, SM32, and SM34. Among them, SM18 (Ergosterol peroxide) was only present in mummified fruits, (Mum_L) and (Mum_T) clusters. SM19 (5,4-dihydroxy-3,6,7-trimethoxy-8C-methylflavone) was present at higher levels in (H-Inj_L) and (Mum_L) clusters compared to the (Asym_L) cluster, and in the (H-Inj_T) and (Mum_T) clusters compared to (Asym_T) cluster. SM20 (5,4-dihydroxy-3,7,8-trimethoxy-6C-methylflavone), SM22 (Apigenin-8-C-β-D-glucopyranoside) and SM25 (Colletoic acid) showed a much higher relative abundance in (Mum_L) and (Mum_T) clusters compared to (Asym_L) and (Asym_T) clusters. However, SM24 (Colletofragarone A1) was found with a high relative abundance in all clusters except for the mummified fruits of both orange cultivars. SM25 (Colletoic acid) showed a much higher relative abundance in (Mum_L) and (Mum_T) clusters compared to (Asym_L) and (Asym_T) clusters. Finally, SM23 (Collectotrichin A) and SM34 (Pyrenocine A) were only present in (Mum_L) and (Mum_T) clusters and were not detected in (Asym_L) and (Asym_T) clusters.

Figures 3A and B show the relative abundance of *Penicillium* metabolites detected in peel and juice of orange fruits. The secondary metabolites produced by *P. digitatum* in the in orange peel were SM35, SM36, SM37, SM38, SM39, SM40, SM42, SM43, SM44, SM45, SM46 and SM47.

**Figure 3.**
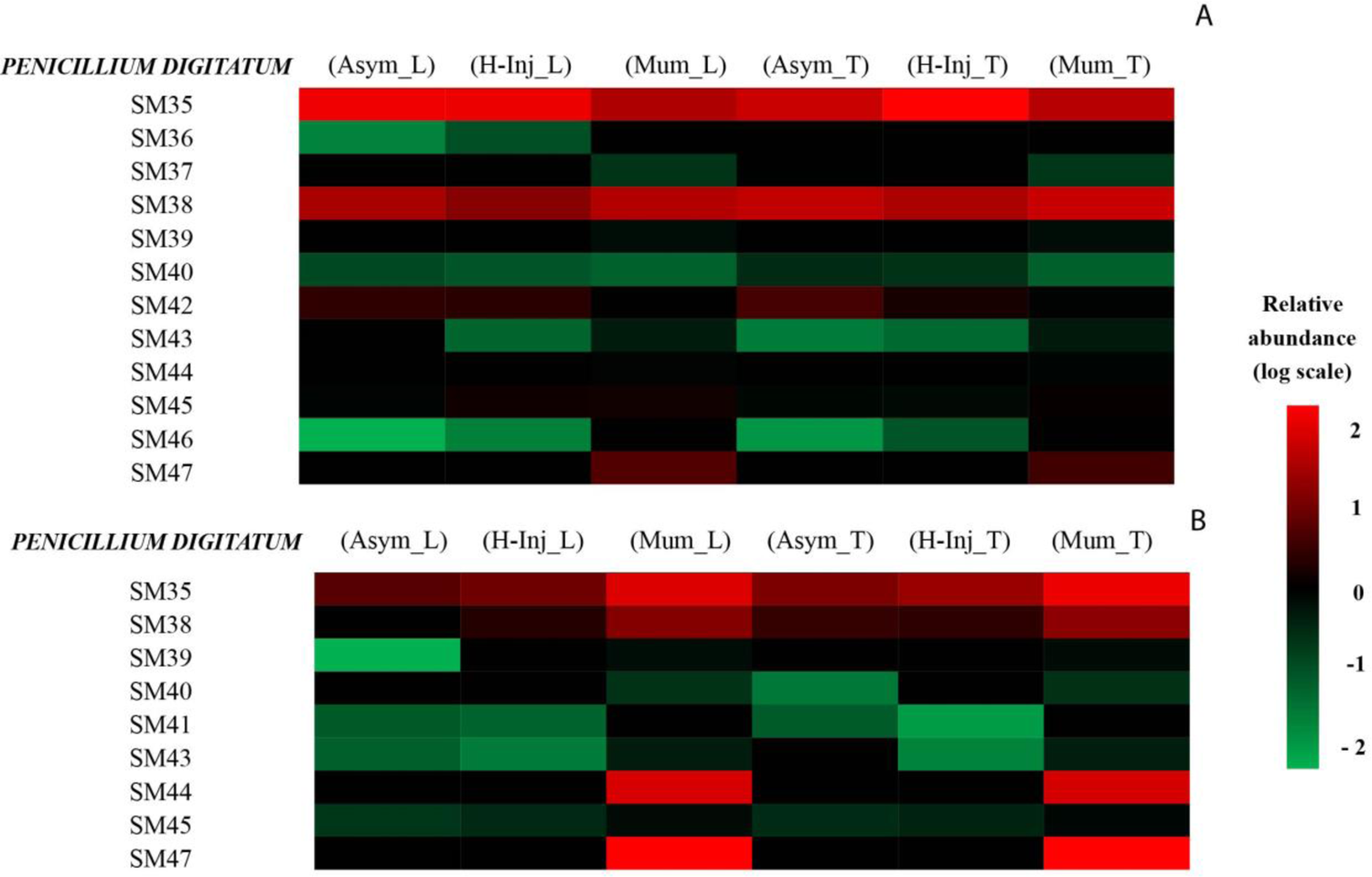
Heat map representing the relative abundance of *Penicillium digitatum* secondary metabo-lites detected in orange peel (A) and orange juices (B). Colors represent the relative abundance (log-arithmic scale) of the secondary metabolites detected: red represents high abundance and green represents low abundance.

However, in the orange peel, only SM35 (Alantrypinone) and SM38 (Atrovenetins) were found with a high relative abundance compared to the other metabolites. Specifically, these secondary metabolites were found with a high relative abundance in all clusters of both orange cultivars. In cluster (H-Inj_L), SM36 (Anacine) was 9-fold more abundant than in cluster (Asym_L). SM37 (Asteltoxin), SM39 (Fungisporin), SM44 (Rubratoxin B) and SM47 (Patulin) were present only in mummified fruits of both orange cultivars, i.e. in (Mum_L) and (Mum_T) clusters. SM42 (Penipacid B) was found with a high relative abundance in all clusters except for the mummified fruits of both orange cultivars. SM43 (Penochalasin K) was detected in hail-injured and mummified fruits, (H-Inj_L) and (Mum_L) clusters, but was almost absent in asymptomatic fruits, cluster (Asym_L). In hail-injured and mummified fruits of ‘Tarocco Tapi’, (H-Inj_T) and (Mum_T) clusters, respectively, this metabolite was about 2- and 63-fold, respectively, more abundant than in asymptomatic fruits, cluster (Asym_T). Finally, SM46 (Solistatin), in hail-injured fruits of both ‘Tarocco Lempso’ and ‘Tarocco Tapi’, clusters (H-Inj_L) and (H-Inj_T), was about 8- and 15-fold, respectively, more abundant than in asymptomatic fruits of both cultivars, clusters (Asym_L) and (Asym_T), respectively.

The secondary metabolites detected in orange juice using the *Penicillium* dataset were SM35, SM38, SM39, SM40, SM41, SM43, SM44, SM45 and SM47. Specifically, SM35 (Alant-rypinone) and SM38 (Atrovenetins) were found with a higher relative abundance compared to the other metabolites. In juice of mummified fruits of both orange cultivars, (Mum_L) and (Mum_T) clusters, the relative abundance of SM35 (Alantrypinone) was approximately 4- and 3-fold higher than in juice of asymptomatic fruits of both cultivars, (Asym_L) and (Asym_T) clusters, respectively. Similarly, the relative abundance of SM38 (Atrovenetins) in mummified fruits of both orange cultivars, (Mum_L) and (Mum_T) clusters, was approximately 4- and 3-fold higher than in asymptomatic fruits, (Asym_L) and (Asym_T) clusters, respectively. In (Mum_L) cluster, SM39 (Fungisporin) showed a relative abundance approximately 355-fold higher than in cluster (Asym_L); it was present in (Mum_T) cluster but was not detected in cluster (Asym_T). SM40 (Lichexanthone) was found in cluster (Mum_L) but not in cluster (Asym_T). Moreover, this metabolite in (Mum_T) showed a relative abundance approximately 13-fold higher than in cluster (Asym_T). SM43 (Penochalasin K), in mummified fruits of ‘Tarocco Lempso’, cluster (Mum_L), showed a relative abundance approximately 11-fold higher than in asymptomatic fruits of the same cultivar, cluster (Asym_L). This metabolite was also detected in mummified fruits of ‘Tarocco Tapi’, cluster (Mum_T), but was not found in asymptomatic fruits of this orange cultivar, cluster (Asym_T). SM44 (Rubratoxin B) and SM47 (Patulin) were found only in (Mum_L) and (Mum_T) clusters. In mummified fruits of the two orange cultivars, clusters (Mum_L) and (Mum_T), SM45 (Serantrypinone) was detected with a relative abundance about 5- and 4-fold higher than in asymptomatic fruits of the respective cultivars, clusters (Asym_L) and (Asym_T).

Principal Component Analysis (PCA) of data was performed. It was based on the secondary metabolites of *A. alternata*, *C. gloeosporioides* and *P. digitatum* (Table 2) identified in peel and juice of asymptomatic, hail-injured and mummified fruits of the two blood orange cultivars, ‘Tarocco Lempso’ and ‘Tarocco Tapi’.

**Table 2.**
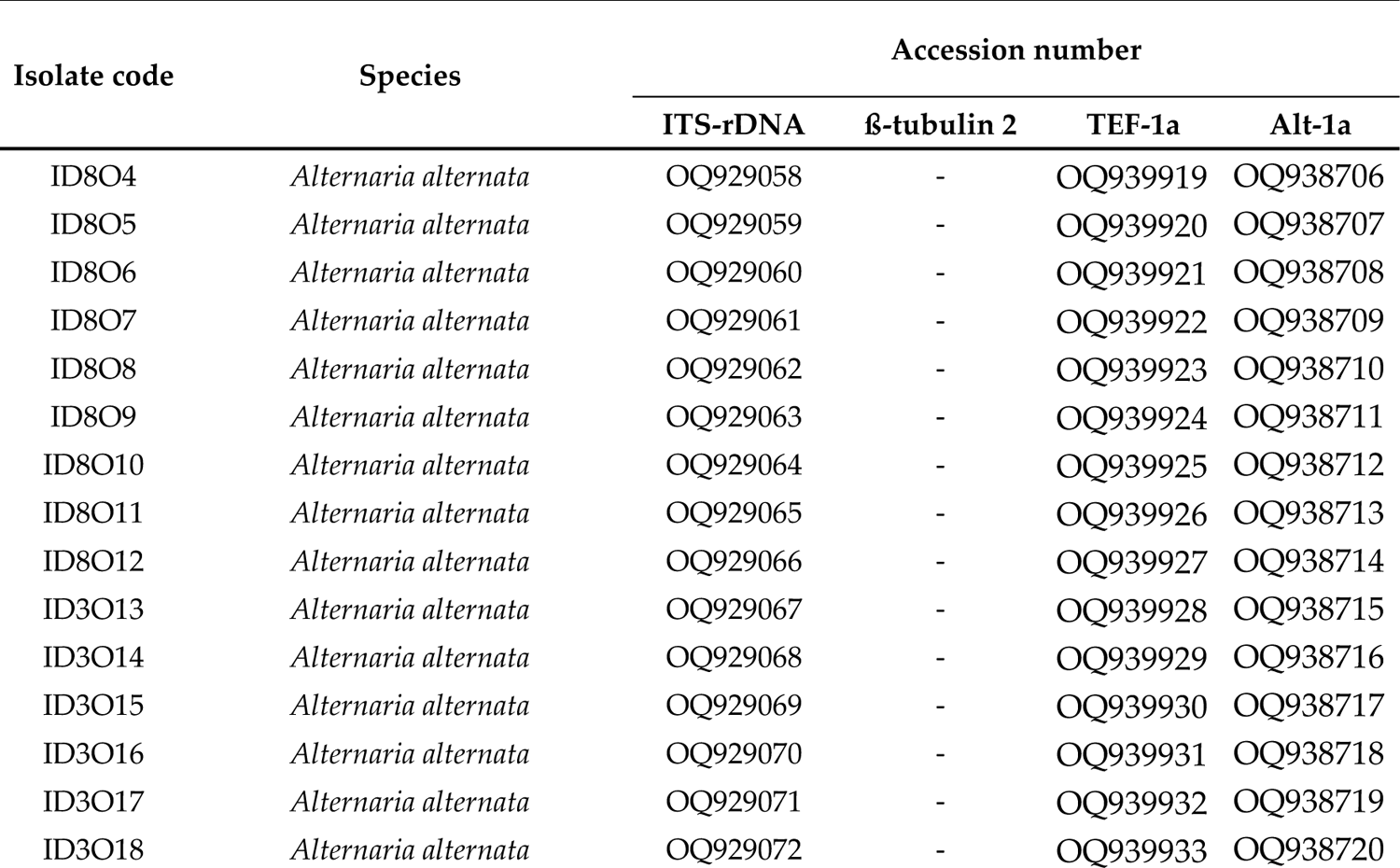
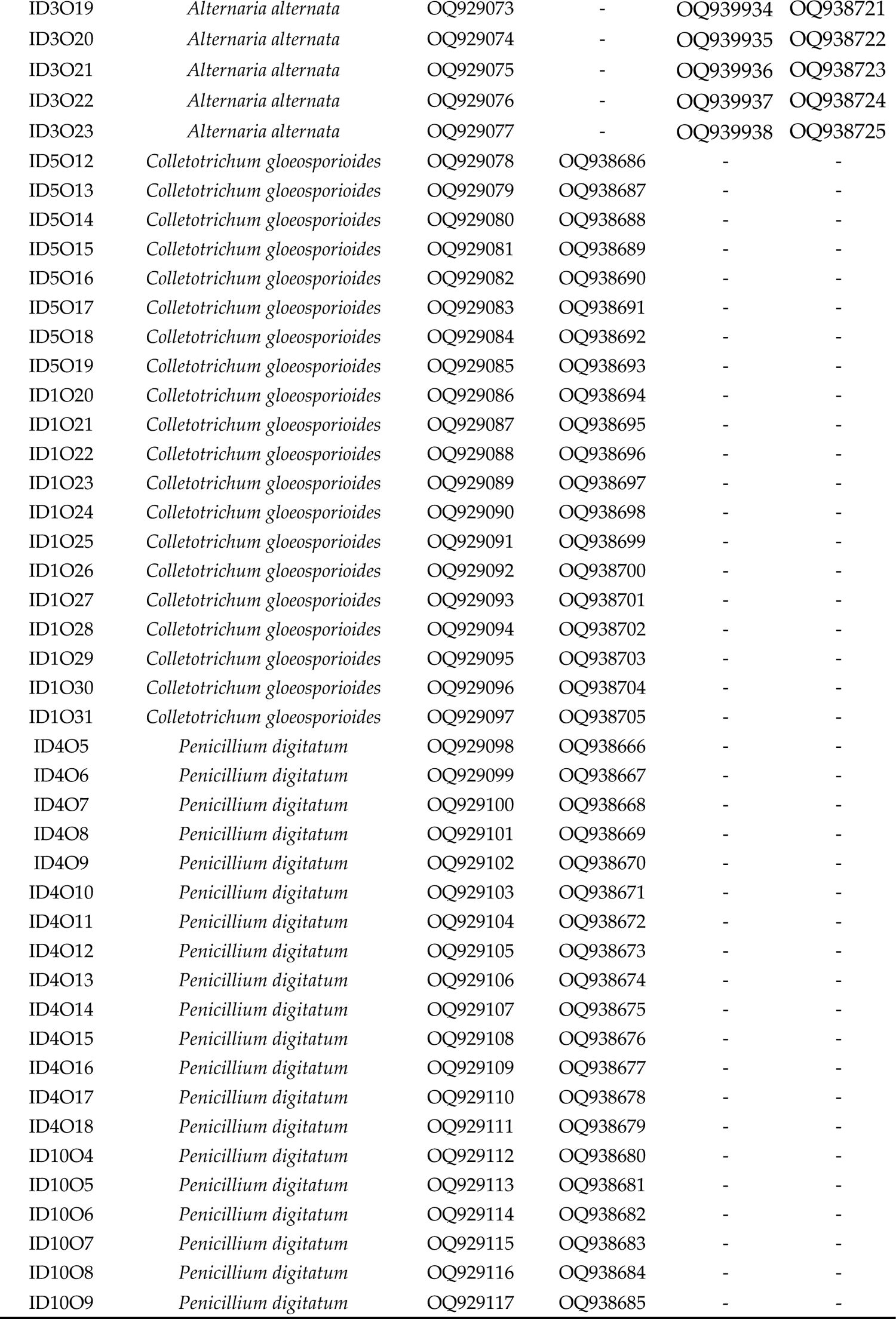
Isolates of *Alternaria*, *Colletotrichum* and *Penicillium* characterized in this study.

Figure 4 shows the clustering of asymptomatic, hail-injured, and mummified orange fruits of the two orange cultivars in the score plot and loading plot, based on secondary metabolites of *Alternaria* detected in the peel. The sum of the principal component values accounted for 69.9% of the total variance, with PC1 representing 47.3% and PC2 representing 22.6% of the total variance.

**Figure 4.**
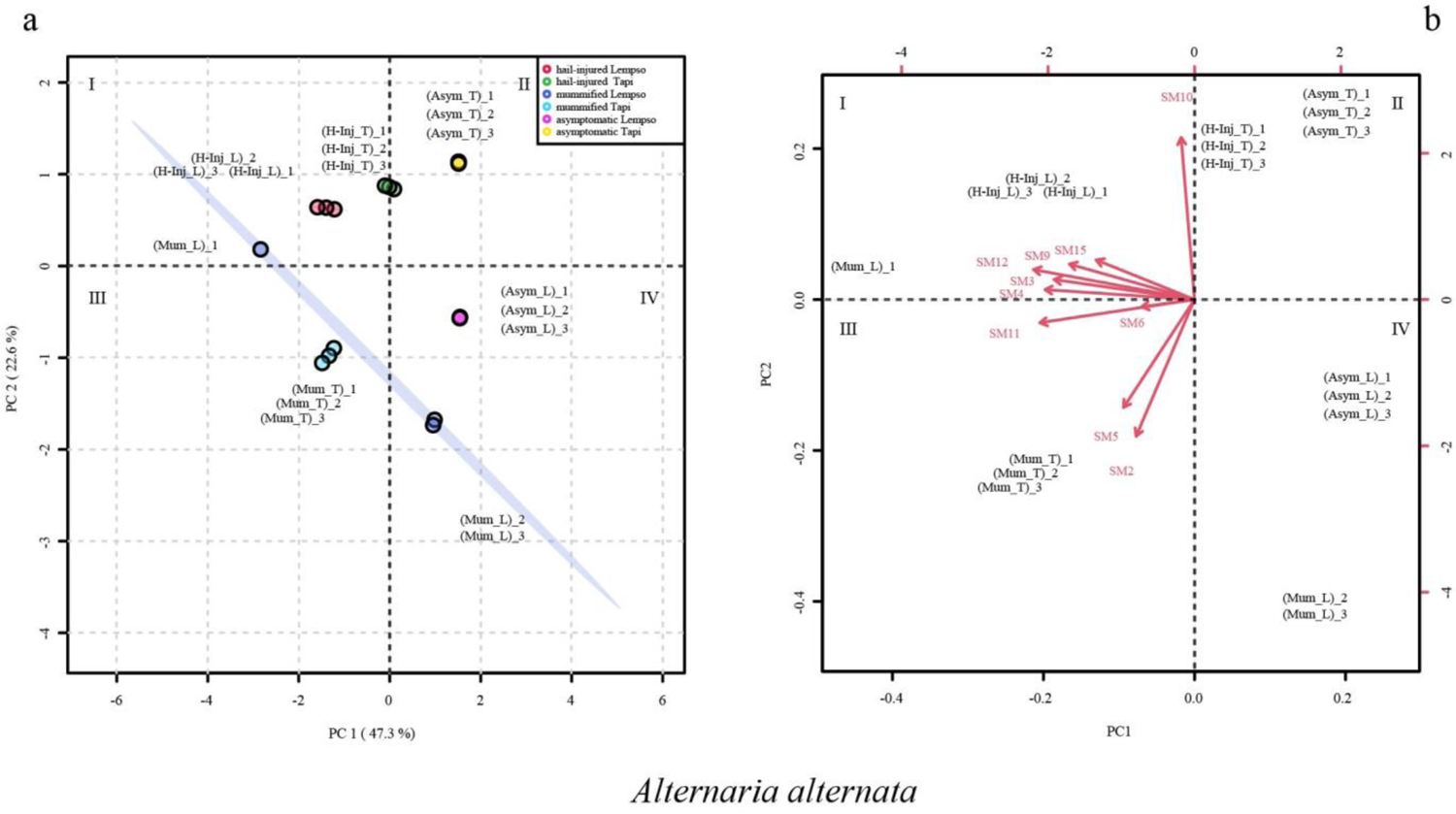
Principal Component Analysis (PCA), scores plot (a) and biplot (b), based on secondary metabolites of *Alternaria alternata* detected in the peel of ‘Tarocco Lempso’ (L) and ‘Tarocco Tapi’ (T) fruits, with three distinct replicates per each type of fruit, asymptomatic (Asym), hail injured (H-Inj) and mummified (Mum).

In the score plot, asymptomatic orange fruits (Asym_T and Asym_L) clustered in quadrants II and IV, while hail-injured fruits (H-Inj_T and H-Inj_L) clustered in quadrant II. Mummified fruits of ‘Tarocco Lempso’ (Mum_L), with the only exception of replicate #1, tended towards quadrant IV. Conversely, mummified fruits of ‘Tarocco Tapi’ (Mum_T) clustered in quadrant III.

In the loading plot, the hail-injured fruits (H-Inj_T and H-Inj_L) clustered in quadrant I, characterized by the secondary metabolites SM3, SM4, SM6, SM9, SM11, SM12 and SM15. By contrast, mummified fruits of ‘Tarocco Tapi’ clustered in quadrant I, characterized by secondary metabolites SM2 and SM5 (Figure 4b).

The score plot and loading plot of Figure 5 illustrate the clustering of the three types of fruits (asymptomatic, hail-injured and mummified) of ‘Tarocco Lempso’ and ‘Tarocco Tapi’ based on secondary metabolites of *Colletotrichum* detected in the peel. The principal component values accounted for 72.2% of the total variance, with PC1 and PC2 accounting for 58.6% and 13.6%, respectively.

**Figure 5.**
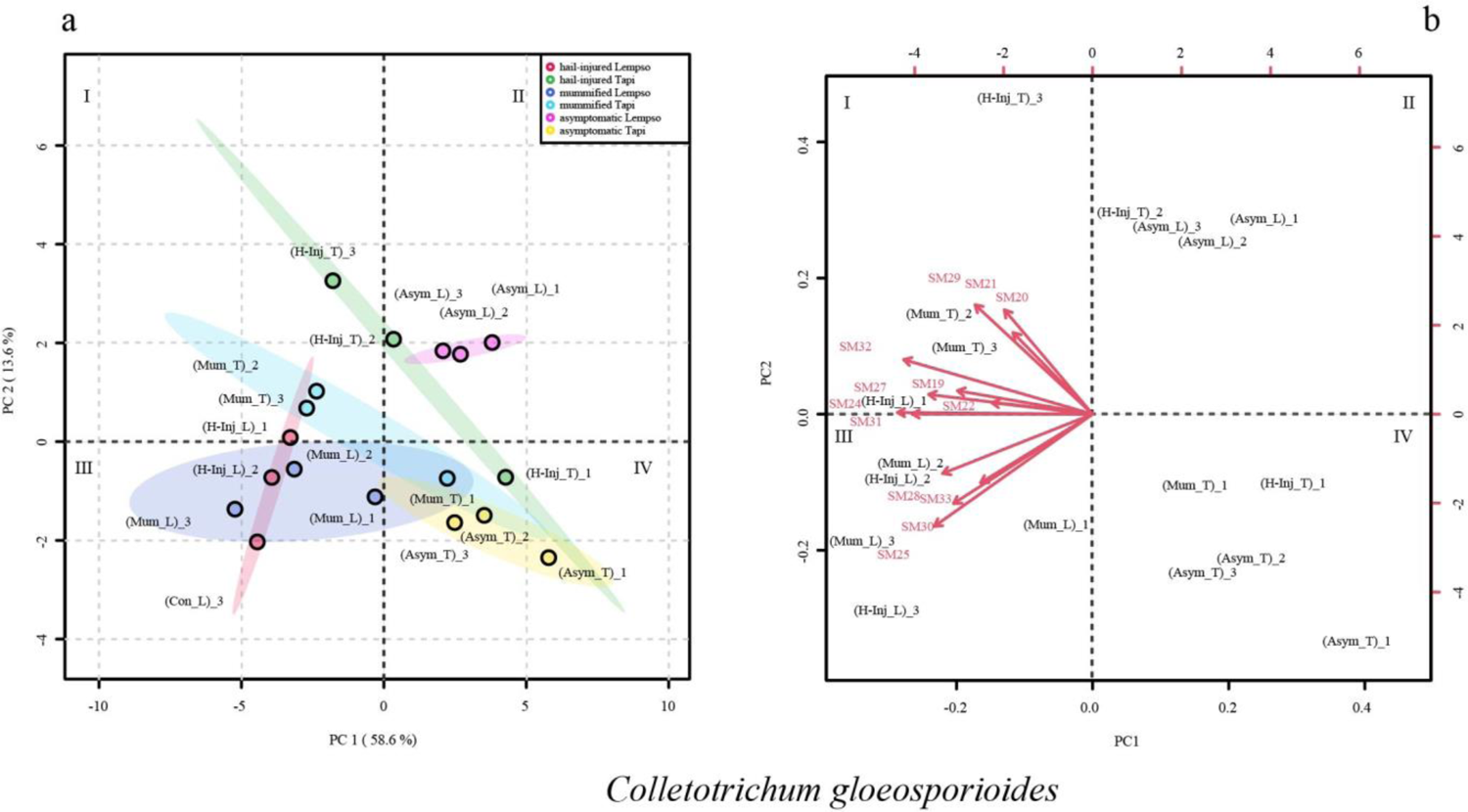
Principal Component Analysis (PCA), scores plot (a) and biplot (b), based on secondary metabolites of *Colletotrichum gloeosporioides* detected in the peel of ‘Tarocco Lempso’ (L) and ‘Tarocco Tapi’ (T) fruits, with three distinct replicates per each type of fruit, asymptomatic (Asym), hail injured (H-Inj) and mummified (Mum).

In the score plot, the asymptomatic fruits of the two orange cultivars (Asym_L and Asym_T) grouped in quadrant II and IV, respectively. The hail-injured fruits of ‘Tarocco Lempso’ (H-Inj_L) clustered in quadrant III, while the hail-injured fruits of ‘Tarocco Tapi’ (H-Inj_T) were spread across quadrants I, II, and IV. Mummified fruits of ‘Tarocco Lempso’ and ‘Tarocco Tapi’ (Mum_L and Mum_T) clustered in quadrants III and I respectively, with the only exception of Mum_T replicate #1, which tended towards quadrant IV.

The loading plot showed that secondary metabolites associated with the peel of mummified fruits of ‘Tarocco Lempso’ (Mum_L) clustering in quadrant I included SM1, SM2, SM3, SM4, SM6, SM9, and SM12, while secondary metabolites.associated with the peel of hail-injured fruits of ‘Tarocco Tapi’ (H-Inj_T), clustering in quadrant I included SM5, SM7, SM8, SM10, SM11, and SM15.

In the loading plot, secondary metabolites SM19, SM22, SM24, SM27, SM28, SM30, SM31, SM32 and SM33 tended towards quadrants I and III, clustering the hail-injured fruits of ‘Tarocco Lempso’ (H-Inj_L) as well as the mummified fruits of both orange cultivars (Mum_L and Mum_T) (Figure 5).

Figure 6 shows the clustering of asymptomatic, hail-injured and mummified fruits of ‘Tarocco Lempso’ and ‘Tarocco Tapi’ on the basis of *Penicillium* secondary metabolites detected in the peel (score plot) and the metabolite trends (loading plot) in different quadrants (I, II, III, IV); the sum of the principal component values accounted for 69.7% of the total variance. PC1 represented 51.1 % and PC2 18.6% of the total variance.

**Figure 6.**
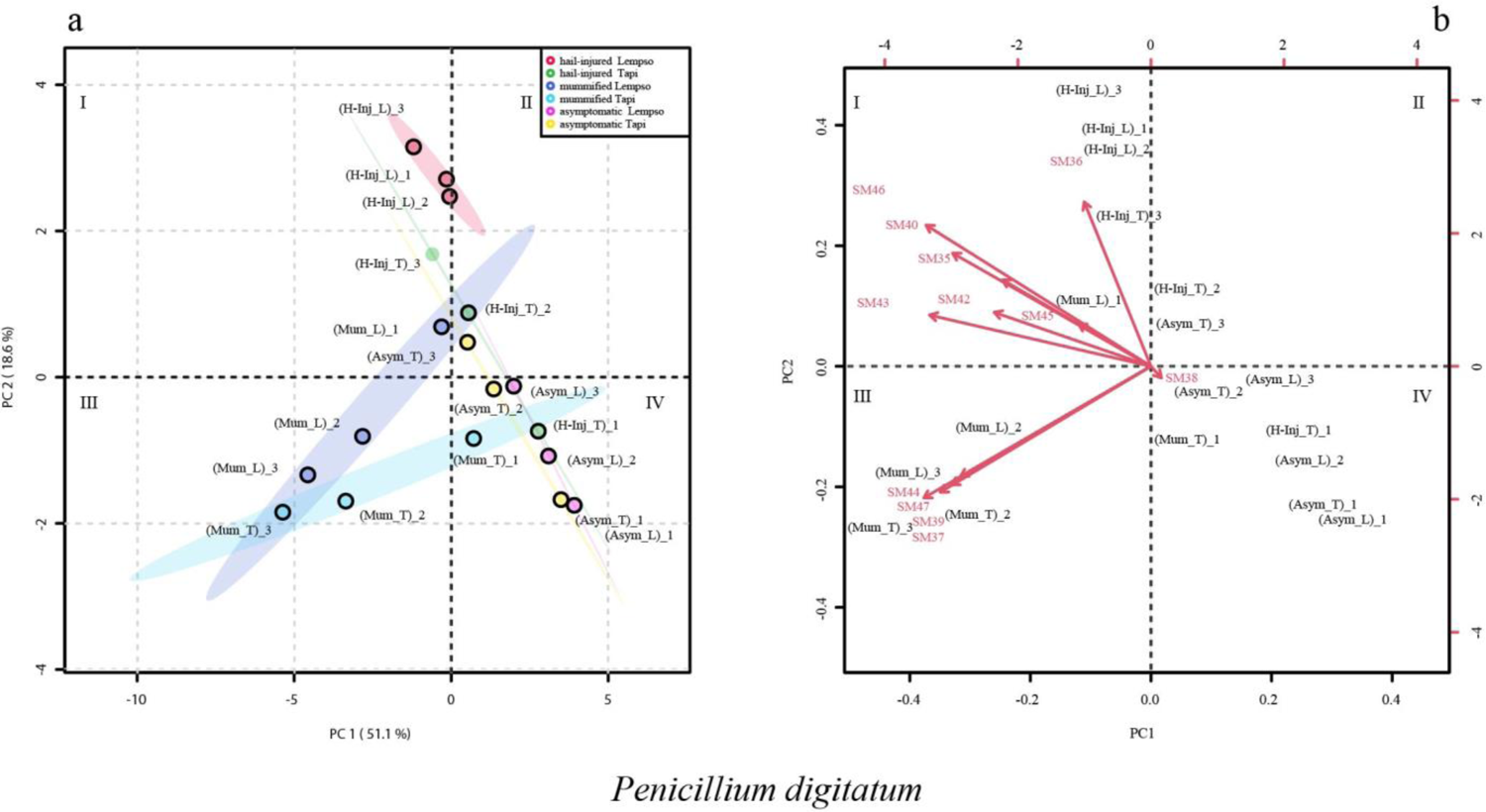
Principal Component Analysis (PCA), scores plot (a) and biplot (b), based on secondary metabolites of *Penicillium digitatum* detected in the peel of ‘Tarocco Lempso’ (L) and ‘Tarocco Tapi’ (T) fruits, with three distinct replicates per each type of fruit, asymptomatic (NoCon), hail injured (Con) and mummified (Mum).

The score plot of *Penicillium* metabolites detected in the peel shows that asymptomatic fruits (Asym_L and Asym_T) clustered in quadrant IV, with the exception of replicate #3, which tended towards quadrant II. Hail-Injured fruitsof ‘Tarocco Lempso’ (H-Inj_L) clustered in quadrant I, while fruits of ‘Tarocco Tapi’ (H-Inj_T) were scattered in quadrants I, II, and IV. The mummified fruits of both cultivars (Mum_T and Mum_L) clustered in quadrant III, with the exception of cluster (Mum_T) replicate # 1, which tended towards quadrant IV.

In the loading plot, secondary metabolites SM35, SM36, SM37, SM39, SM40, SM42, SM43, SM44, SM45, SM46 and SM47 tended towards quadrants I and III, clustering the samples of peel of hail-injured fruits of ‘Tarocco Lempso’ (H-Inj_L), replicate #3 of hail-injured fruits of ‘Tarocco Tapi’ (H_Inj_T_3) and mummified fruits of both orange cultivars (Mum_L and Mum_T) (Figure 6).

Figure 7 shows the clustering of asymptomatic, hail-injured and mummified orange fruits of ‘Tarocco Lempso’ and ‘Tarocco Tapi’ based on *Alternaria* metabolites (score plot) detected in the juice and the metabolite trends (loading plot) in different quadrants (I, II, III, IV). The sum of the principal 12 component values reached 76.3% of the total variance. PC1 and PC2 accounted for 48.9 and 27.4% of the total variance, respectively.

**Figure 7.**
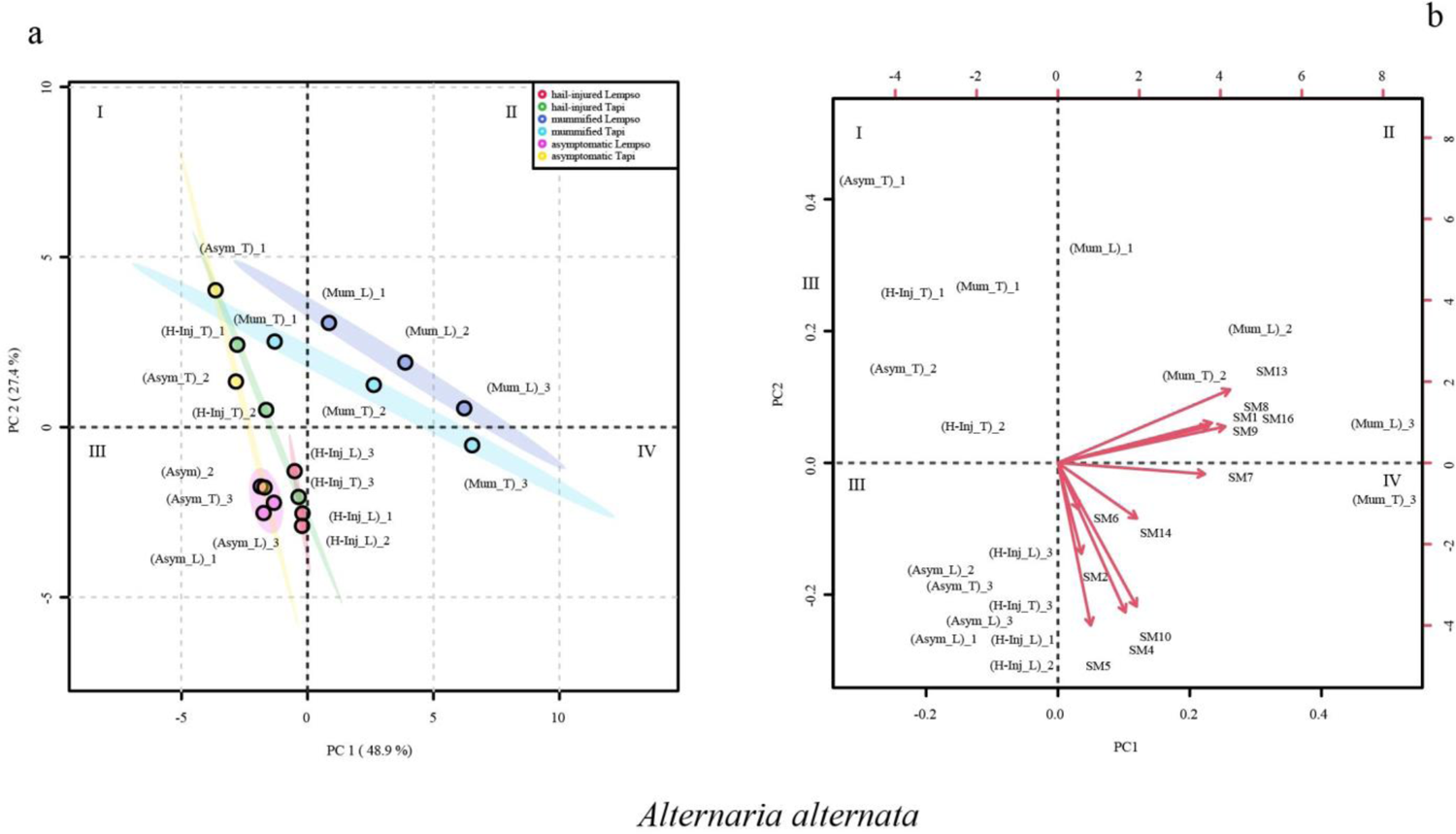
Principal Component Analysis (PCA), scores plot (a) and biplot (b), based on secondary metabolites of *Alternaria alternata* detected in the juice of ‘Tarocco Lempso’ (L) and ‘Tarocco Tapi’ (T) fruits, with three distinct replicates per each type of fruit, asymptomatic (Asym), hail injured (H-Inj) and mummified (Mum).

In the score plot of *Alternaria alternata*, asymptomatic fruits of the two orange cultivars (Asym_T and Asym_L) clustered in quadrants I and III, respectively, with the only exception of replicate #3 of asymptomatic fruits of ‘Tarocco Tapi’ (Asym_T_3), which tended towards quadrant III. Hail-injured fruits of ‘Tarocco Lempso’ (H-Inj_L) clustered in quadrant III, while hail-injured fruits of ‘Tarocco Tapi’ (H-Inj_T) clustered in quadrant I, with the only exception of replicate #3 (H-Inj_T_3) that tended towards quadrant III. Mummified fruits of ‘Tarocco Lempso’ (Mum_L) clustered in quadrant II, while mummified fruits of ‘Tarocco Tapi’ (Mum_T) clustered in quadrants I, II and IV.

In the loading plot, the secondary metabolites SM1, SM7, SM8, SM13 and SM16 tended towards quadrant II, clustering the mummified fruits of ‘Tarocco Lempso’ (Mum_L and Mum_T), and the metabolites SM2 SM4, SM5, SM6, SM10 and SM14 clustered the hail-injured fruits of both orange cultivars (H-Inj_L and Mum_T) (Figure 7).

Figure 8 shows the clustering of asymptomatic, hail-injured and mummified fruits of ‘Tarocco Lempso’ and ‘Tarocco Tapi’, based on *Colletotrichum* secondary metabolites detected in the juice (score plot) and the metabolite trends (loading plot) in different quadrants (I, II, III, IV). The sum of the principal 14 component values accounted for 70.3% of the total variance. PC1 and PC2 accounted for 50.9 and 19.4% of the total variance, respectively.

**Figure 8.**
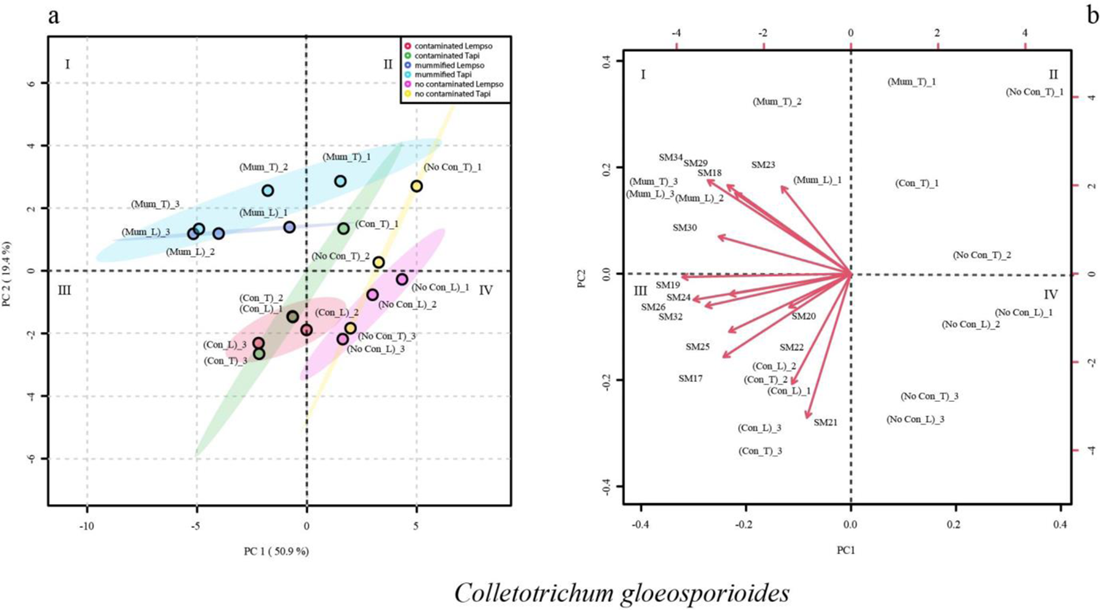
Principal Component Analysis (PCA), scores plot (a) and biplot (b), based on secondary metabolites of *Colletotrichum gloeosporioides* detected in the juice of ‘Tarocco Lempso’ (L) and ‘Tarocco Tapi’ (T) fruits, with three distinct replicates per each type of fruit, asymptomatic (Asym), hail injured (H-Inj) and mummified (Mum).

In the score plot of *Colletotrichum gloeosporioides*, asymptomatic fruits of both orange cultivars (Asym_T and Asym_L) clustered in quadrants II and IV, respectively, with the only exception of replicate #3 of ‘Tarocco Tapi’ (Asym_T_3), which tended towards quadrant IV. Hail-injured fruits of both orange cultivars (H-Inj_L and H-Inj_T) clustered in quadrant III, with the only exception of replicate #1 of hail-injured ‘Tarocco Tapi’ (H-Inj_T_1), which tended towards quadrant II. Mummified fruits of both orange cultivars (Mum_L and Mum_T) clustered in quadrant I, with the only exception of replicate #1 of mummified ‘Tarocco Tapi’ (Mum_T_1), which tended towards quadrant II. In the loading plot, the secondary metabolites, SM18, SM19, SM23, SM29, SM30 and SM34, tended towards quadrant I, clustering the mummified fruits of both orange cultivars (Mum_L and Mum_T), while the secondary metabolites SM17, SM20, SM24, SM25, SM26 and SM32 tended towards quadrant III, clustering the hail-injured fruits of both orange cultivars (H-Inj_L and H-Inj_T) (Figure 8).

Figure 9 shows the clustering of asymptomatic, hail-injured and mummified fruits of ‘Tarocco Lempso’ and ‘Tarocco Tapi’, based on *Penicillium secondary* metabolites detected in the juice (score plot) and the metabolite trends (loading plot) in different quadrants (I, II, III, IV). The sum of the principal nine component values accounted for 77.2% of the total variance. PC1 and PC2 accounted for 60.2 and 17% of the total variance, respectively. In the score plot of *Penicillium digitatum*, asymptomatic fruits of the two orange cultivars (Asym_T and Asym_L) clustered in quadrants I and III. Hail-injured fruits of both orange cultivars (H-Inj_L and H-Inj_T) clustered in quadrants I and II, while mummified fruits of the two cultivars (Mum_L and Mum_T) clustered in quadrants III and IV. In the loading plot, the secondary metabolites SM38, SM39, SM41 and SM45 tended towards quadrant II, clustering the hail-injured fruits of both ‘Tarocco Lempso’ and ‘Tarocco Tapi’ (H-Inj_L and H-Inj_T), while the secondary metabolites SM40 and SM47 tended towards quadrant IV, clustering the mummified fruits of both orange cultivars (Mum_L and Mum_T) (Figure 9).

**Figure 9.**
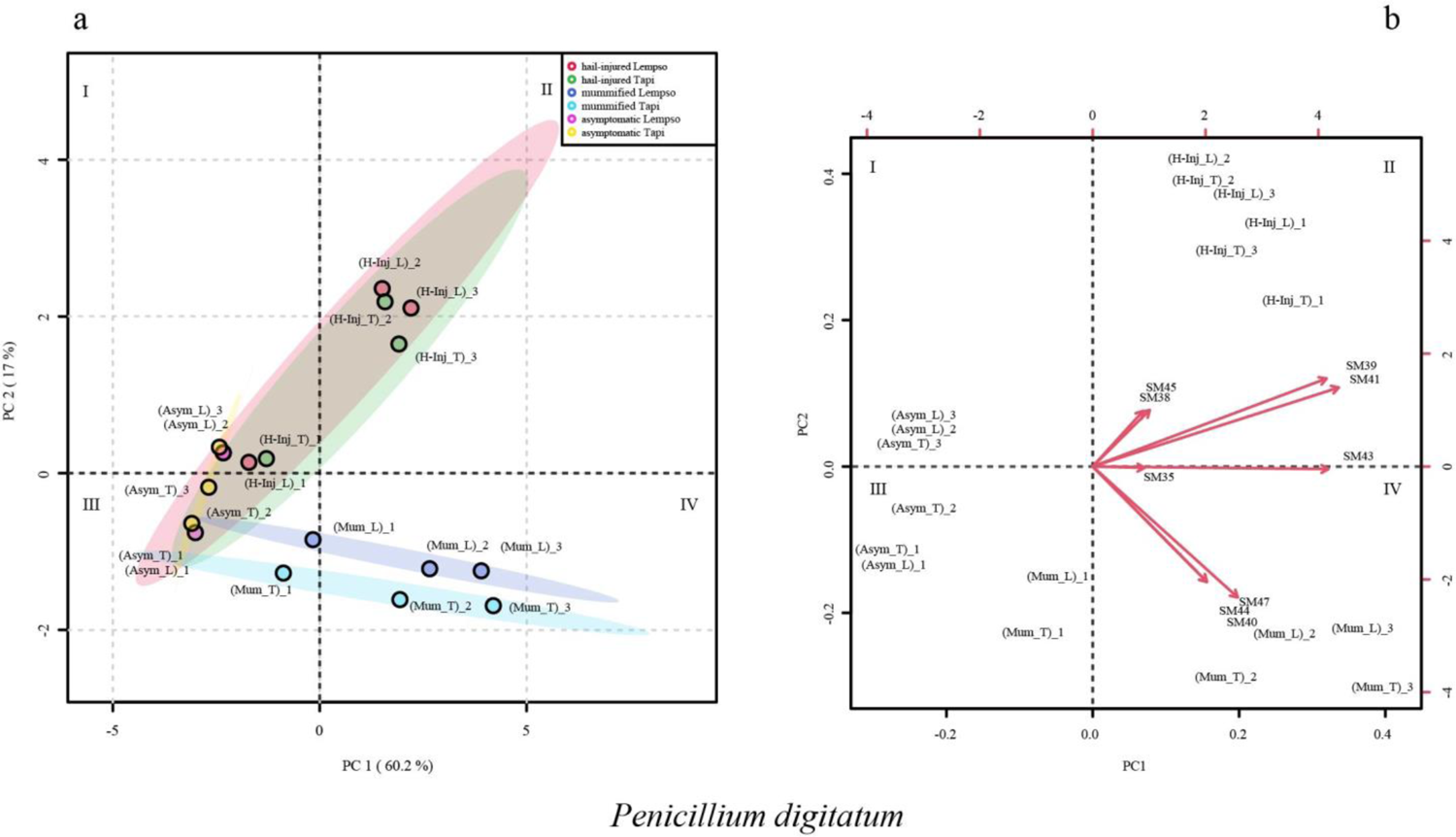
Principal Component Analysis (PCA), scores plot (a) and biplot (b), based on secondary metabolites of *Penicillium digitatum* detected in the juice of ‘Tarocco Lempso’ (L) and ‘Tarocco Tapi’ (T) fruits, with three distinct replicates per each type of fruit, asymptomatic (Asym), hail injured (H-Inj) and mummified (Mum).

To summarize the results reported above for each fungal species, the metabolic profiles of peel extracts of Tarocco Lempso’ and ‘Tarocco Tapi’ and the metabolic profiles of the three types of fruits overlapped only in part, as shown by Ven diagrams (Figure 10). In particular, the secondary metabolites Colletotrilactam (SM29) and Patulin (SM47) were produced exclusively in the peel of mummified fruits of both cultivars. In contrast, all clusters shared the secondary metabolite Alternethanoxin A (SM6), 5,4-dihydroxy-3,7,8-trimethoxy-6C-methylflavone (SM20), apigenin-8-C-β-D-glucopyranoside (SM22), alantrypinone (SM35) and Atrovenetins (SM38). AF-toxins II (SM3) and Penipacid B (SM42) were found in the peel of hail-injured fruits of both cultivars, while Colletomelleins B (SM27) and Fusarentin 6,7-dimethyl ether (SM 31) were found exclusively in the peel of hail-injured fruits of ‘Tarocco Lempso’ and ‘Tarocco Tapi’, respectively. Finally, Serant-rypinone (SM45) was detected in the peel of both hail-injured and asymptomatic fruits but only in fruits of ‘Tarocco Lempso’.

**Figure 10.**
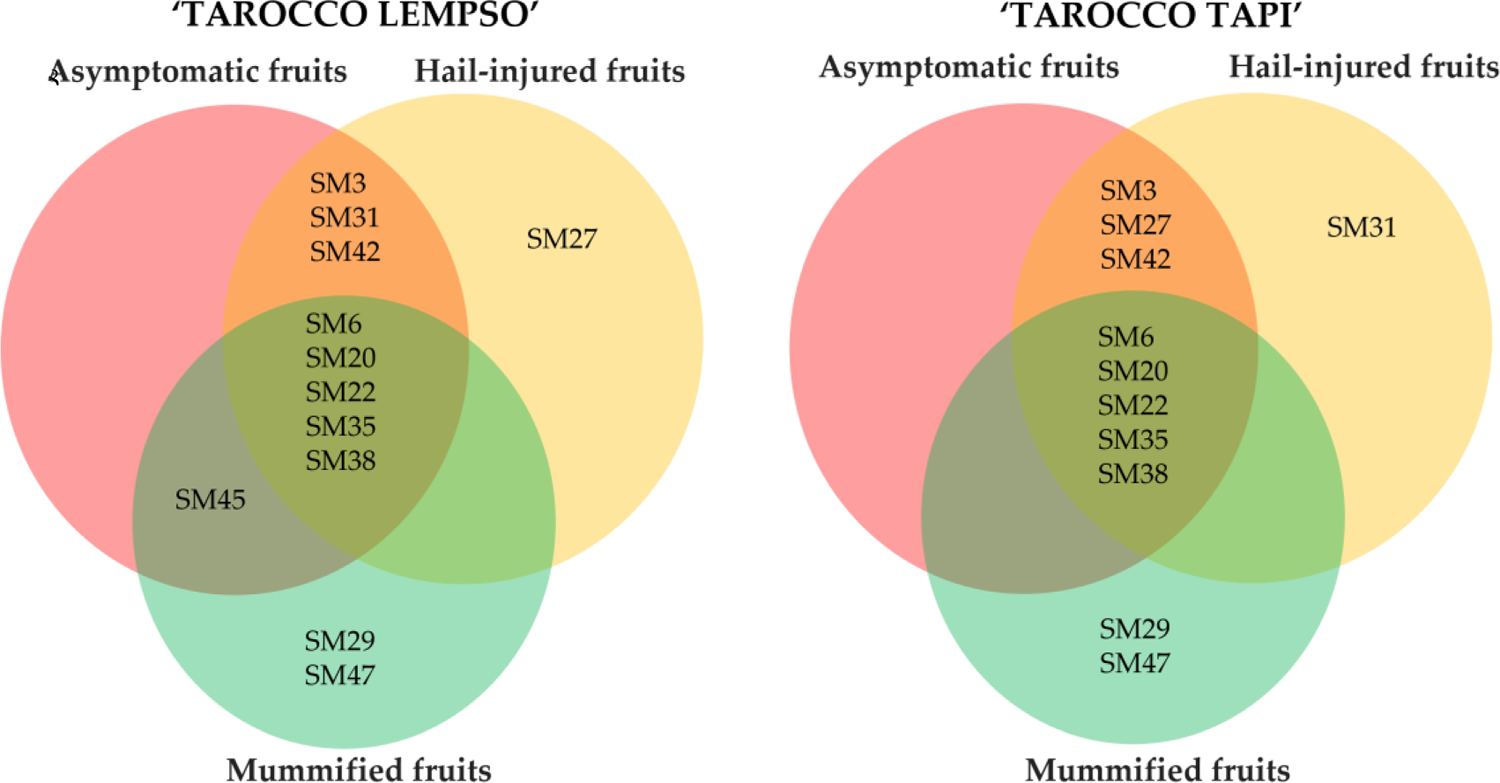
Venn diagrams showing the distribution of secondary metabolites in peel extracts of hail-injured, mummified and asymptomatic fruits of ‘Tarocco Lempso’ and ‘Tarocco Tapi’ cultivars.

The metabolic profiles of the juice from mummified fruits of both ‘Tarocco Lempso’ and ‘Tarocco Tapi’ cultivars (Figure 11) had in common several secondary metabolites, including AAL-toxin TE2 (SM2), Alternethanoxin A (SM6), ergosterol peroxide (SM18), apigenin-8-C-β-D-glucopyranoside (SM22), Colletoic acid (SM25), Rubratoxin B (SM44) and Patulin (SM47). However, while SM2, SM18, SM22, SM25, SM44 and SM47 were found exclusively in the profile of mummified fruits, SM6 was found in the profile of all types of fruits. Similarly, Alantrypinone (SM35) was found in all types of fruits but only in those of ‘Tarocco Lempso’, while 5,4-dihydroxy-3,7,8-trimethoxy-6C-methylflavone (SM20) and Atrovenetins (SM38) were found in all types of fruits of ‘Tarocco Tapi’ and only in hail-injured and mummified fruits of ‘Tarocco Lempso’. The metabolic profiles of asymptomatic and hail-injured fruits of both orange cultivars shared Porriolide (SM14) and Colletofragarone A1 (SM24), while (+) - (3R,4S)-cis-4-hydroxy-6-deoxyscytalone was detected exclusively in the profile of asymptomatic fruits of ‘Tarocco Tapi’.

**Figure 11.**
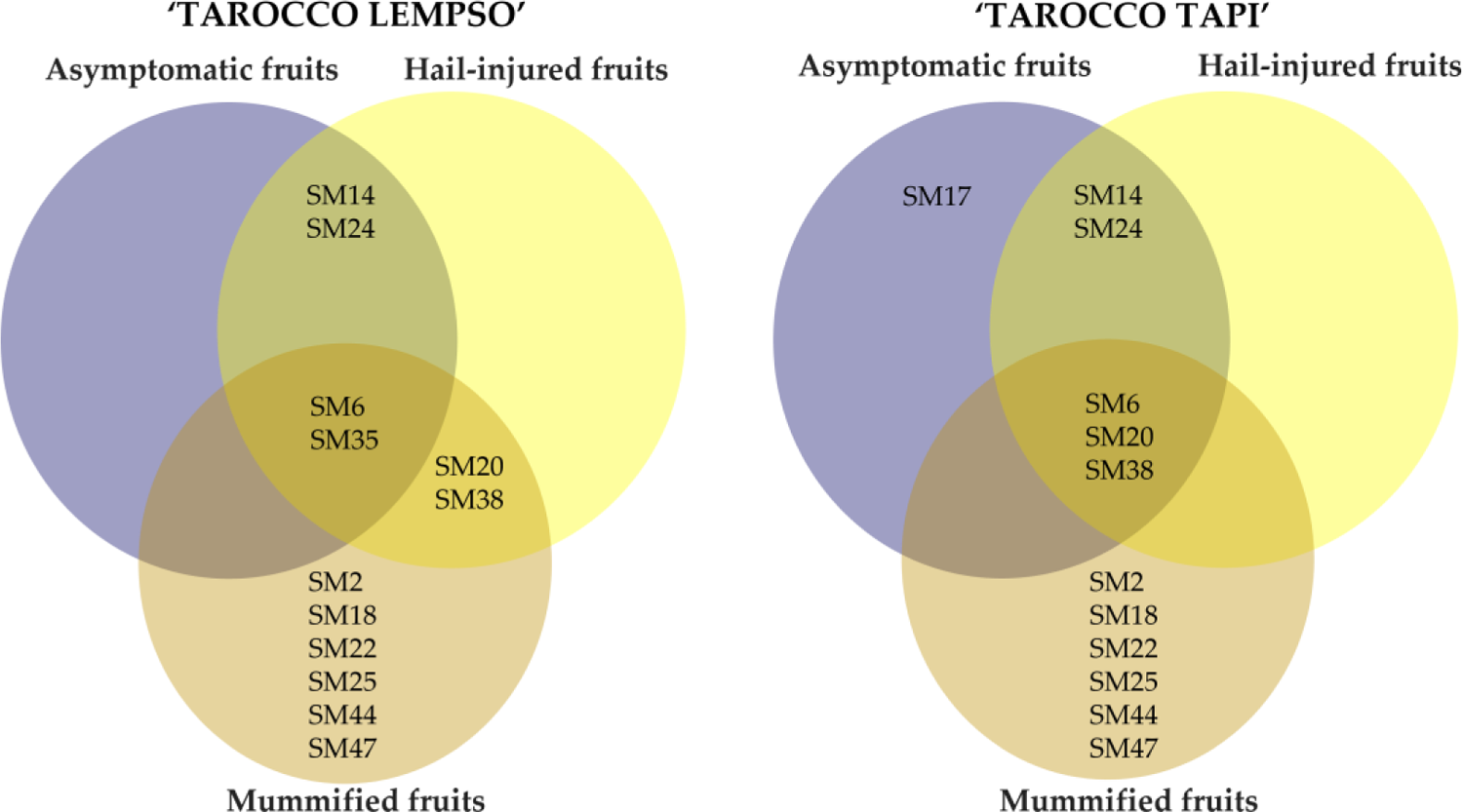
Venn diagrams showing the distribution of secondary metabolites in juice of hail-injured, mummified and asymptomatic fruits of ‘Tarocco Lempso’ and ‘Tarocco Tapi’ cultivars.

## 3. Discussion

In this study, isolations with a conventional microbiological method from the fruit peel of two blood orange cultivars, ‘Tarocco Lempso’ and ‘Tarocco Tapi’, sampled in commercial citrus orchards of a typical production area of eastern Sicily, yielded three species of fungi, *A. alternata*, *C. gloeosporioides* and *P. italicum*. Fruits of each cultivar were separated into three distinct groups on the basis of external symptoms, i.e. fruits with the rind injured by the hail, mummified fruits and asymptomatic fruits, with no visible lesions of the rind. *Penicillium italicum* was the prevalent fungal species recovered from both mummified and hail-injured fruits of both orange cultivars suggesting this fungus had a preminent role in fruit mummification and confirming its nature of wound parasite. Both *C. gloeosporioides* and *A. alternata* were recovered from necrotic lesions of hail-injured fruits of both orange cultivars in a significant higher frequency than from the intact rind of asymptomatic fruits. These fungi are ubiquitous in citrus orchards and normally reside in the peel of citrus fruits as epiphytes, endophytes or latent pathogens. Very probably, in fruits injured by hail their populations increased because they colonized the rind lesions as opportunistic necrotrophic pathogens. A possible explanation of both the failure in isolating *A. alternata* and the low proportion of successful isolations of *C. gloeosporioides* from mummified fruits is that *P. digitatum* exerted an antagonistic activity against the other two fungal species, based on the competition for the substrate or the production of fungitoxic secondary metabolites by this fungus. *Alternaria alternata*, *C. gloeosporioides* and *P. digitatum* are responsible for some of the most common and severe pre- and post-harvest diseases of citrus fruits worldwide [1,13,23]. Moreover, they belong to three among the most known genera of mycotoxigenic fungi [58,124]. Overall, *Alternaria* species produce more than 70 mycotoxins sensu lato [94] and more than 100 secondary metabolites of *Colletotrichum* species have been reported so far [103]. *Penicillium* species produce a wide range of mycotoxins of concern for human and animal health. Among mycotoxins occurring in foods and feeds contaminated by *Penicillium* species, the most important are ochratoxin A and patulin, which are regulated in a number of countries, and to a lesser extent cyclopiazonic acid [124]. The identification of culturable fungi recovered from the peel of ‘Tarocco Lempso’ and ‘Tarocco Tapi’ fruits oriented the design and development of *Alternaria*, *Colletotrichum* and *Penicillium* metabolite databases, that in this study were essential tools for the exploitation of UHPLC–Q-TOF-MS analysis method. Very recently, this innovative analytical approach has been successfully used for identifying the secondary metabolites produced by diverse *Colletotrichum* species both in vitro and on olives infected by anthracnose [125]. Actually, in the present study the combined application of UHPLC–Q-TOF-MS and databases of fungal metabolites proved to be very effective in identifying the secondary metabolites released by *Alternaria*, *Colletotrichum* and *Penicillium* in the peel and juice of fruits of ‘Tarocco Lempso’ and ‘Tarocco Tapi’. Overall, as many as 47 diverse fungal secondary metabolites were identified in the peel and juice of fruits of these two blood orange cultivars. Of these metabolites, 16 were produced by A. alternata, 18 by *C. gloeosporioides* and 13 by *P. digitatum*. According with the scientific literature all these metabolites are functionally active and most of them show diverse types of activity. However, not all bioactive secondary metabolites produced by *A. alternata*, *C. gloeosporioides* and *P. italicum* detected in the peel and juice of citrus fruits in this study can be regarded as mycotoxins sensu strictu [28] and [126].

In particular, as for the metabolites of *Alternaria*, Alternariol monomethyl ether (AME) is a mycotoxin that can cause kidney damage and immunosuppression in humans and animals [102,127,128]. AME was also reported to induce mitochondrial apoptosis in human colon carcinoma cells [90]. Alternethanoxin A displayed an in vitro inhibitory activity of the growth of cancer cells [123]. Aurasperone-C is known to cause nervous system dysfunctions [129]. Curvularin, which in this study was found only in mummified fruits, was reported to be a carcinogenic and immunotoxic substance [130]. Altenusin (ALN) was reported to be a multifunctional metabolite, with antibacterial, antifungal and antiparasitic activity [131]. The amount of this metabolite in orange fruits varied considerably. In juice of hail-injured fruits, its relative abundance was three to six fold higher than in asymptomatic fruits while in the juice of mummified fruits its relative abundance was even 63- to 251-fold higher than in asymptomatic fruits. AAL-toxins were demonstrated to have phytotoxic, cytotoxic and genotoxic activity [88,132]. AAL-toxin TE2 was found with a high relative abundance in the juice of mummified orange fruits of both cultivars. This metabolite belongs to the AAL-toxins, a group of mycotoxins produced by *A. alternata* structurally related to Fuminosins produced by *Fusarium* species. AAL-toxins were reported as contaminats of various crops and as host specific phytoxins (HSTs) [94,133]. In the *A. alternata*/tomato pathosystem, they have a key role as pathogenicity factors [88,134,135]. AF-toxin II is a mixture of three stereoisomers reported to be phytotoxic on strawberry and pear fruits [135]. In this study, AF-toxin II is reported for the first time on citrus fruits infected by A. alternata. This metabolite was detected only in the peel of asymptomatic and hail-injured fruits of both orange cultivars examined. Maculosin was proposed as a potential eco-friendly agrochemical with herbicidal activity [94], while Porriolide was shown to exert a significant antifungal activity [136]. In this study, Alternethanoxin A was found with a high relative abundance in all types of orange fruits, including asymptomatic, hail-injured and mummified fruits. In the original description of this metabolite isolated from A. sonchi cultures, the Authors envisaged the possibility of using it as a mycoerbicide in view of its broad-spectrum phytotoxic activity [137].

As for *Colletotrichum* metabolites identified in this study, Colletofragarone A1, Colletoic acid and Pyrenocine A were reported to have phytotoxic and, in a few cases, cytotoxic properties [138,139]. Other compounds such as 5,4-dihydroxy-3,7,8-trimethoxy-6C-methylflavone and Apigenin-8-C-β-D-glucopyranoside, which in this study were found with a high relative abundance in all types of orange fruits examined, are reported in the literature as bioactive metabolites [101,103]. Apigenin, in particular, was demonstrated to have a broad-spectrum anticancer activity [103,140]. However, no one of these *Colletotrichum* metabolites has been so far reported as a mycotoxin of concern for human and animal health.

As for *Penicillium*, it is known this fungus produces a wide range of secondary bioactive metabolites, including metabolites with cytotoxic activity such as Penicillic acid, Pat-ulin and Ochratoxin A [124,141]. Patulin is a mycotoxin that has been found as contaminant in various types of fruits, including oranges [142]. It has been identified as a potential hazard for human health and food safety [18]. Several studies evidenced Patulin can cause adverse health effects in humans, including cytotoxicity, genotoxicity, and immunotoxicity [42,143]. The European Union and the United States Food and Drug Administration have set maximum limits for Patulin in food products, including oranges [26].

In this study, consistently with the results of isolations, high levels of Patulin were detected in mummified fruits of both orange cultivars, in particular in juice. Rubratoxin B was another secondary metabolite found with a high relative abundance in the juice of mummified orange fruits of both ‘Tarocco Lempso’ and ‘Tarocco Tapi’. It is produced by several species of *Penicillium* [144]. Rubratoxin B is a mycotoxin exhibiting a range of acute and chronic toxic effects, including hepatotoxicity, nephrotoxicity, splenotoxicity and genotoxicity. It is commonly found as contaminant in cereals, such as rice and wheat, and poses a potential health risk to humans and animals [126,145]. Lichexanthone is an additional secondary metabolite found in a considerable amount in the juice of mummified orange fruits. This is a well-known metabolite of lichens and was also found in many filamentous fungi, including endophytic *Penicillium* and *Trichoderma* species [146]. It has been reported to possess antimicrobial and antitumoral activity. Other secondary metabolites of Penicillium, recovered in orange fruits, included Alantrypinone, a metabolite with herbicidal activity [147], Serantrypinone a metabolite with insecticidal activity [148], Atrovenetin known for its antioxidant effects [114], Asteltoxin and Penochalasin K, both with proven antifungal activity [114,149]. Other Penicillium secondary metabolites with antimicrobial and antifungal activity such as Fungisporin, Palitantin and Penipacid B were found in orange fruits. These metabolites were detected in both mummified and hail injured fruits and their presence seems correlated with the prevalence of this fungus over the other fungal species as inferred from the high proportion of Penicillium isolates recovered from these two types of fruits. By analogy with other plant pathogenic fungal species [150], it can be speculated that the ability of *P. digitatum* to produce several metabolites, which show allelopathic, antifungal, and antibacterial activity, might enhance the ecological fitness of this fungus. Moreover, this feature could explain the ability of *P. digitatum* to exclude or inhibit other less competitive species, such as *A. alternata* and *C. gloeosporioides*, from mummified fruits or to overgrow them during isolation of axenic cultures.

The PCA of the interactions among driving factors shaping the profile of secondary metabolites in orange fruits showed the profiles of mummified fruits were distinct from the profiles of either asymptomatic or hail-injured fruits. It is noteworthy the mycotoxin Patulin was a characteristic component of the *Penicillium* metabolite profile of both the peel and juice of mummified fruits of both orange cultivars examined while the other *Penicillium*-related mycotoxin Rubratoxin B was detected with a high relative abundance only in the juice of this type of fruits. Moreover, the *Penicillium* metabolite profile from the peel of hail-injured fruits was distinct from the profile of asymptomatic fruits and was characterized by the presence of secondary metabolites, such as Alantrypinone, Anacine, Lichexanthone, Penipacid B, Serantrypinone and Solistatine, whose toxic activity for humans and animals has been so far little investigated.

As for *C. gloeosporioides*, the metabolic profiles in both the peel and juice of hail-injured fruits of ‘Tarocco Lempso’ and ‘Tarocco Tapi’ grouped separately from the profiles of the corresponding asymptomatic fruits, suggesting an interaction between hail injuries and the biology of this fungus which is a common resident in the fruit peel but also a latent pathogen.

As for *A. alternata*, all the secondary metabolites identified in this study were recovered from mummified and hail-injured fruits. Only one of them, AME, has been reported as a mycotoxin of concern for human health produced by Alternaria species associated to citrus [24]. Little is known about the toxicological potential of the other Alternaria-related metabolites recovered from orange fruits in this study or the risk of fruit contamination.

Most of the studies aimed to characterize the biological activity of these metabolites or the ability of fungal isolates to produce them were performed in vitro. In general, to evaluate the risk of food and feed contamination, it is noteworthy that overall 26 out of 47 metabolites, identified in this study, were recovered from both the fruit peel and the juice (6, 9 and 8 metabolites of *A. alternata*, *C. gloeosporioides* and *P. digitatum*, respectively), 12 exclusively from the peel (4 per each of the three fungal species) and 12 exclusively from the juice (6 and 5 of *A. alternata* and *C. gloeosporioides*, respectively, one of *P. digitatum*). Consequently, the risk of contamination concerns both fresh fruits and processed products such as juices, marmelade and citrus pulp, the last one used largely as a feed for cattle.

## 4. Materials and Methods

### 4.1 Orange fruit sampling

On 24^th^ January 2022, mature fruits of two blood orange cultivars, ‘Tarocco Lempso’ and ‘Tarocco Tapi’, were collected in three 20-years-old commercial orchards cultivated with organic farming methods, in three diverse districts of the province of Catania (eastern Sicily). Around five months before the sampling a hailstorm had beaten the orchards and since then trees had not been treated with fungicides. Three distinct types of fruits were selected per each cultivar: fruits injured by hail (fruits with necrotic lesions covering around 25% of the rind surface), mummified fruits still on the tree canopy, and asymptomatic fruits (fruits with no visible rind injury) (Figure 12).

**Figure 12.**
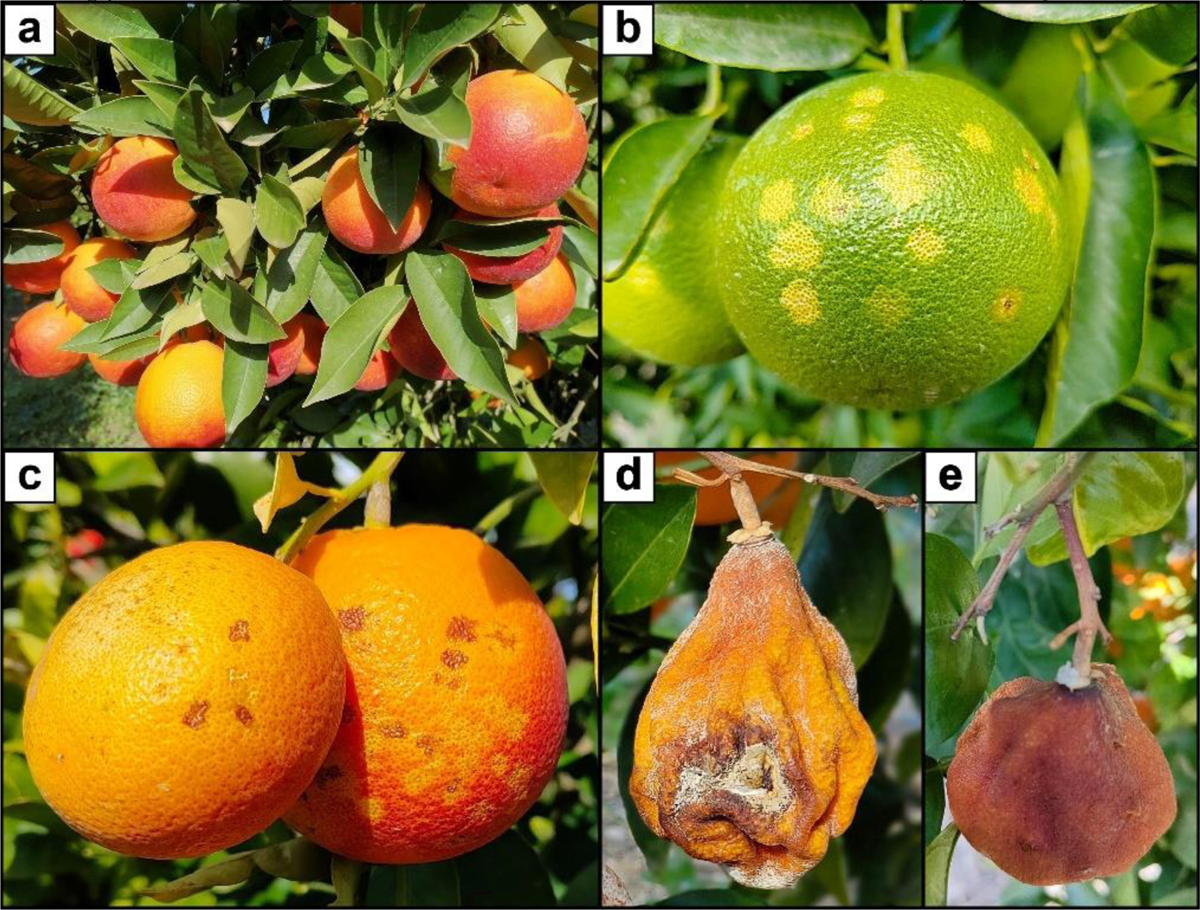
**(a)** Asymptomatic mature fruits of ‘Tarocco Lempo’; **(b)** Rind blemishes (oleocellosis) on a green fruit in autumn, few days after the hail-storm; **(c)** Blemished (hail-injured) mature fruits of ‘Tarocco Lempso’; **(d-e)** Mummified fruit of ‘Tarocco Lempso’.

Overall, in each orchard 60 fruits of each orange cultivar were collected, five fruits of each type (hail-injured, mummified and asymptomatic) were picked up randomly from four randomly selected trees. Overall, 60 fruits of the same type and cultivar were pooled together and three replicates, each of 20 fruits, for each composite sample were processed and analyzed separately (Table 4). The sampling design is displayed in Table 4.

**Table 3.** Secondary metabolites of *Alternaria alternata*, *Colletotrichum gloeosporioides* and *Penicillium digitatum* identified in this study in the peel and juice of blood orange fruits.

| Secondary metabolites | ID code <sup>a</sup> | Matrices <sup>b</sup> | Chemical class | Reference <sup>c</sup> |
| --- | --- | --- | --- | --- |
| AAL-toxin TB2 | SM1 | juice | Mycotoxins<br>(Cyclic peptides) | [88] |
| AAL-toxin TE2 | SM2 | peel and juice | Mycotoxins | [88] |
| AF-toxin II | SM3 | peel | Mycotoxins<br>(Fumonisin) | [89] |
| Altenusin (ALN) | SM4 | peel and juice | Benzoxazole | [58] |
| Alternariol monomethyl ether (AME) | SM5 | peel and juice | Mycotoxins | [90] |
| Alternethanoxin A | SM6 | peel and juice | Ketone | [90] |
| Altersolanol L | SM7 | juice | Derivatives<br>of anthraquinone | [92] |
| Aurasperone C | SM8 | juice | Prenylated benzophenones | [93] |
| Curvularin | SM9 | peel and juice | Macrocyclic lactones | [94] |
| Dihydroaltersolanol | SM10 | peel and juice | Cyclohexenones | [95] |
| Erythroglaucon | SM11 | peel | Anthraquinones | [88] |
| Macrosporin A | SM12 | peel | Macrolides | [58] |
| Maculosin | SM13 | juice | Tetramic acids | [96] |
| Porriolide | SM14 | juice | Furanones | [97] |
| Porritoxinol | SM15 | peel | Phthalides | [98] |
| Tentoxin (TEN) | SM16 | juice | Cyclic<br>tetrapeptides | [58] |
| (+) - (3R,4S)-cis-4-hydroxy-6-deoxyscytalone | SM17 | juice | Tetramic acids | [99] |
| Ergosterol peroxide | SM18 | juice | Steroids | [100] |
| 5,4-dihydroxy-3,6,7-trimethoxy-8C-methylflavone | SM19 | peel and juice | Flavones | [101] |
| 5,4-dihydroxy-3,7,8-trimethoxy-6C-methylflavone | SM20 | peel and juice | Flavones | [101] |
| Alternariol-5-Me ether | SM21 | peel and juice | Coumarins | [102] |
| Apigenin-8-C-β-D-glucopyranoside | SM22 | peel and juice | Flavonoid<br>glycosides | [103] |
| Collectotrichin A | SM23 | juice | Cyclopeptides | [104] |
| Colletofragarone A1 | SM24 | peel and juice | Sesquiterpenoids | [105] |
| Colleteoic acid | SM25 | peel and juice | Diketopiperazines | [106] |
| Colletoketol | SM26 | juice | Macrolides | [101] |
| Colletomelleins B | SM27 | peel | Quinones | [107] |
| Colletonoic acid | SM28 | peel | Carboxylic acids | [108] |
| Colletotrilactam A | SM29 | peel and juice | Lactams | [109] |
| Colletotrilactam D | SM30 | peel and juice | Lactams | [109] |
| Fusarentin 6,7-dimethyl ether | SM31 | peel | Coumarins | [110] |
| Novae-zelandin A | SM32 | peel and juice | Phenazines | [111] |
| Phthalide | SM33 | peel | Cyclic anhydrides | [110] |
| Pyrenocine A | SM34 | juice | Pyrenocines | [112] |
| Alantrypinone | SM35 | peel and juice | Quinazolinones | [88] |
| Anacine | SM36 | peel | Quinazolinones | [113] |
| Asteltoxin | SM37 | peel | Anthraquinones | [114] |
| Atrovenetins | SM38 | peel and juice | Xanthoness | [114] |
| Fungisporin | SM39 | peel and juice | Cyclohexenones | [115] |
| Lichexanthone | SM40 | peel and juice | Xanthoness | [116] |
| Palitantin | SM41 | juice | Flavones | [117] |
| Penipacid B | SM42 | peel | Diketopiperazines | [118] |
| Penochalasin K | SM43 | peel and juice | Macrolides | [119] |
| Rubratoxin B | SM44 | juice | Mycotoxins | [120] |
| Serantrypinone | SM45 | peel and juice | Quinones | [121] |
| Solistatin | SM46 | peel | Lactones | [122] |
| Patulin | SM47 | peel and juice | Mycotoxins<br>(Furanocoumarins) | [46] |

<sup>a</sup> SM1-SM16: *Alternaria alternata*; SM17-SM34: *Colletotrichum gloeosporioides*; SM35-SM47 *Penicillium digitatum*. 208
<sup>b</sup> Matrices on which the secondary metabolites were identified in this study. 210
<sup>c</sup> Original reports of the secondary metabolites produced by *Alternaria*, *Colletotrichum* and *Penicillium* spp. 211

**Table 4.** Sampling and processing design for the determination of metabolite profile of fruits of two blood orange cultivars.

| Cluster | Number of fruits | Cultivar |
| --- | --- | --- |
| Asymptomatic fruits (Asym_L) | 60<sup>a</sup> | Tarocco Lempso |
| Hail-injured fruits (H-Inj_L) | 60 |  |
| Mummified_fruits (Mum_L) | 60 |  |
| Asymptomatic fruits (Asym_T) | 60 | Tarocco Tapi |
| Hail-injured fruits (H-Inj_T) | 60 |  |
| Mummified_fruits (Mum_T) | 60 |  |

<sup>a</sup> Three replicates, each of 20 fruits.

Fruit samples were kept into plastic bags and immediately shipped to the Laboratory of Food Toxicology at the Department of Preventive, Medicine, Nutrition and Food Science Area of the University of Valencia, Spain. Samples were stored at 4 °C until isolation of fungi from the rind. Subsequently, the juice was extracted and secondary metabolites from the peel and juice were analyzed separately.

### 4.2 Isolation of fungi

Isolations were performed from the fruit peel according with Riolo et al., [17], with few modifications. Fruits were washed with tap water, surface sterilised with a sodium hypochlorite solution (10%) for 1 min, immersed in 70% ethanol for 30 s and rinsed in sterile distilled water (s.d.w.). After disinfection, 15 peel fragments (5×5 mm) per each fruit were taken with a scalpel (**Figure 13**). In fruits damaged by hail, fragments were taken from necrotic lesions of the rind, while in mummified and asymptomatic fruits fragments were taken randomly from the entire surface of the fruit. The fragments were blotted dry with sterile filter paper and placed in Petri dishes (three Petri dishes per fruit) on Potato Dextrose Agar (PDA; Oxoid Ltd., Basingstoke, UK) amended with 150 μg/mL streptomycin and incubated for 2 days at 25°C, in the dark. Pure cultures of each fungal isolates were obtained by single-conidium subcultures.

**Figure 13.**
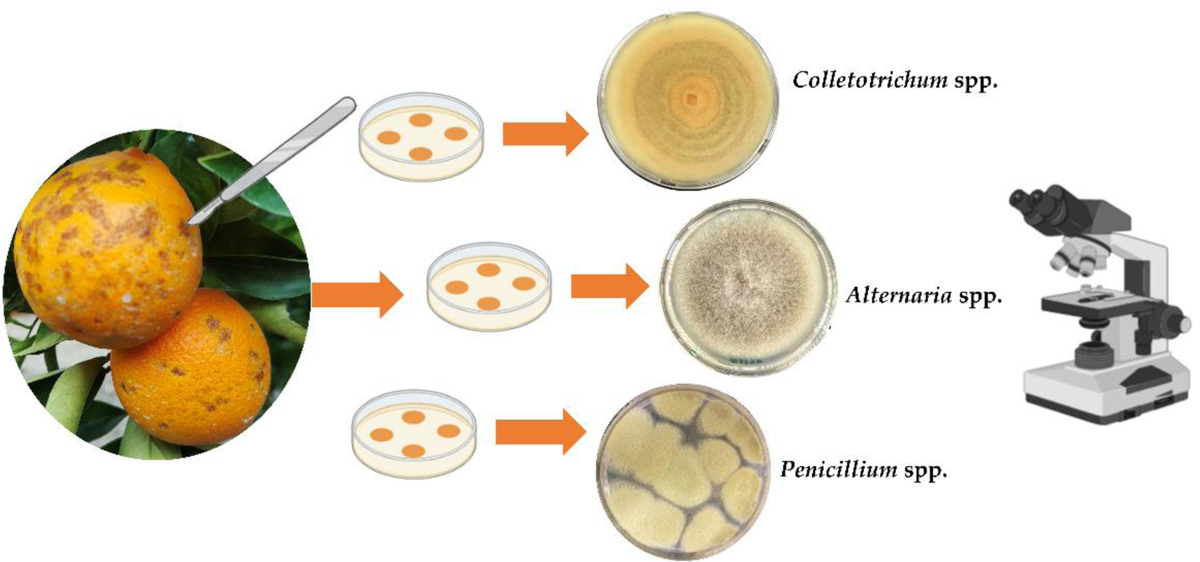
Diagrammatic representation of isolation and morphological identification at genus level of isolates obtained from orange fruits.

### 4.3 Morphological characterization of isolates

The macroscopic characteristics (colour, margin, diameter and texture) of colonies were examined according to Pryor and Michailides [151], whereas microscopic features (conidium and conidiophore branching morphology) according to Simmons [152]and Agrios [153] (Figure 13). Macro- and microscopic identification were used to separate the isolates by genus.

### 4.4 Molecular characterization of isolates

Genomic DNA was extracted from single-conidium isolates using the procedure described in Riolo et al. 2021. Overall, 290 selected fungal isolates of three distinct genera obtained from orange fruits were examined. The internal transcribed spacer (ITS) region of the ribosomal DNA was amplified and sequenced with primers ITS1/ITS4 [154], part of the translation elongation factor 1 alpha gene (tef1) was sequenced and amplified with primers EF1-728F/EF1-986R [155], the β-tubulin gene (tub2) was sequenced and amplified with Bt2a and Bt2b [156] and Alt-1a allergen protein [157] was sequenced and amplified with Alt-for/Alt-rev primers (ITS, EF-1α and Alt-1a allergen for *Alternaria*, ITS and TUB2 for both *Colletotrichum* and *Penicillium*).

The PCR amplifications were performed on a GeneAmp PCR System 9700 (Applied Biosystems, Monza-Brianza, Italy). All PCRs were performed by using the Taq DNA polymerase recombinant (Invitrogen™, Carlsbad, 254 CA, USA) and carried out in a total volume of 25 µL containing the following: PCR Buffer (1X), dNTP mix (0.2 mM), MgCl2 (1.5 mM), forward and reverse primers (0.5 µM each), Taq DNA Polymerase (1 U), and 1 µL of genomic DNA [34]. Amplified products were analyzed by electrophoresis and single bands of the expected size were purified with the QIAquick PCR Purification Kit (Qiagen, Hilden, Germany) and sequenced with both forward and reverse primers by Colección Española de Cultivos tipo Parc Científic (Universitat de València (Valencia, España).

The identification was performed by comparing the sequences with other known sequences in the literature and present on the NCBI (https://blast.ncbi.nlm.nih.gov/Blast) and Mycobank (http://www.mycobank.org)/CBS-KNAW (Westerdijk Fungal Biodiversity Institute) (http://www.westerdijkinstitute.nl).

### 4.5 Preparation of samples for the analyses and extraction of orange juice

Peel was manually separated from albedo using a knife and the fresh orange juice was extracted from peeled and pressed oranges using a domestic squeezer. The peel and orange juice of each fruit were separated and individually transferred into conical polytetrafluoroethyl (PTFE) centrifuge tubes (15 mL). Before the extraction process, all the samples were stored in polyethylene tubes maintained at −80°C.

### 4.6 Extraction of fungal secondary metabolites from orange peel and juice

The extraction of fungal secondary metabolites of orange peel was carried out according with the method of Widiantini et al. [158], partially modified. At first, 5 mL of methanol per each gram of orange peel were added into a conical polytetrafluoroethylene (PTFE) centrifuge tube (15 ml); then was shaken for 12 hours at room temperature with an orbital laboratory shaker. The supernatant was collected and filtered through a 13-mm/0.22-μm nylon syringe filter (Membrane Solutions) in a 2 mL Amber Vial for chromatography analysis and 20μL of this solution was injected for UHPLC– Q-TOF-MS analysis.

The extraction of fungal metabolites of orange juice was carried out according to the method of El Jai et al., [159] slightly modified. At first, 10 mL of orange juice and 20 mL of ethyl acetate (extraction solvent) were added in a conical polytetrafluoroethyl (PTFE) centrifuge tube (50 ml) and the suspension was mixed for 1 min, then it was centrifuged for 10 min at 4000 rpm at 5°C, using an Eppendorf centrifuge 5810R (Eppendorf, Hamburg, Germany). The organic phase at the top of the tube was recovered and placed in another PTFE centrifuge tube (15 mL) and saved. The total collected organic phase was evaporated to dryness under a nitrogen stream using a TurboVap LV evaporator (Zymark, Hopkinton, MA, USA). The dry extract was reconstituted with 1 mL of methanol and filtered through a 13-mm/0.22-μm nylon syringe filter (Membrane Solutions) in a 2mL Amber Vial for chromatography analysis.

### 4.7 Analysis of fungal secondary metabolites by UHPLC– Q-TOF-MS

Orange peel and orange juices extracts were injected for UHPLC– Q-TOF-MS analysis. All the analyses were performed in triplicate (n = 3). The samples were analyzed using an UHPLC device (1290 Infinity II LC, Agilent Technologies) composed of an automatic sampler, a binary pump and a vacuum degasser, coupled with a quadrupole time of flight mass spectrometer (Agilent 6546 LC/Q-TOF) operating in positive ionization mode (Agilent Technologies, Santa Clara, CA, USA), was used for chromatographic analysis. 5 µL of sample was injected. Complete analysis was performed in 25 minutes. Chromatographic separation was performed with an Agilent Zorbax RRHD SB-C18, 2.1 mm x 50 mm, 1.8 µm column. Mobile phase A was composed of Milli-Q water and acetonitrile was used for mobile phase B (both phases were acidified with 0.1% formic acid), with gradient elution, as follows: 0 min, 2% B; 22 min 95% B; 25 min, 5% B. The column was equilibrated for 3 min before every analysis. The flow rate was 0.4 mL/min. Dual AJS ESI source conditions were as follows: gas temperature: 325 C; gas flow: 10 L/min; nebulizer pressure: 40 psig; sheath gas temperature: 295 C; sheath gas flow: 12 L/min; capillary voltage: 4000 V; nozzle voltage: 500 V; Fragmentor: 120 V; skimmer: 70 V; product ion scan range: 100–1500 Da; MS scan rate: 5 spectra/s; MS/MS scan rate: 3 spectra/s; maximum precursors per cycle: 2; and collision energy: 10, 20, 40 eV Untargeted LC/Q-TOF based metabolomics approach were used to identify the different secondary metabolites of fungi growing on orange peel and orange juice. Integration, data elaboration, and identification of metabolites were managed using MassHunter Qualitative Analysis Software B.08.00 and library PCDL Manager B.08.00 (Dopazo et al., 2021).

### 4.8 Statistical analysis

The analytical data obtained by UHPLC– Q-TOF-MS were log10 transformed before statistical analysis. Relationships among two different cultivar (‘Tarocco Lempso’ and ‘Ta-rocco Tapi’) x *Alternaria*, *Colletotrichum* and *Penicillium* species combinations were analyzed using Pearson’s correlation coefficient analysis. All the above statistical analyses were performed using RStudio v.1.2.5 (R). MetaboAnalyst 5.0 software [160] was used for principal component analysis (PCA) using log10 transformed data. The features included were log-transformed and mean-centred.

## 5. Conclusions

Diverse secondary metabolites produced by three among the most common fungi responsible for pre- and post-harvest decays of citrus fruits, were extracted directly from fruits sampled in commercial citrus orchards. They were identified using the modern analytical method of UHPLC–Q-TOF-MS. Many of these metabolites were isolated for the first time from citrus. The medium- and long-term objectives of this study were to evaluate the risk of contamination by mycotoxins and to provide basic information for setting maximum tolerance limits of mycotoxins in fresh citrus fruits and their derivatives. We are aware the latter is a more complex process and need a multidisciplinary approach, as both the dietary habits of targeted populations and the additive effects of toxins from different food sources have to be taken into consideration. Regarding the risk of mycotoxin contamination of citrus fruits, an interesting finding is that with the analytical method we used a consistent level of secondary metabolites was detected even in asymptomatic fruits. Moreover, the metabolic profile of the peel extract of asymptomatic fruits did not differ substantially from the profile of the peel extract of hail-damaged fruits. This is consistent with the results of isolations, which confimed that *Alternaria*, *Colletotrichum* and *Penicillium* reside in the peel of asymptomatic fruits as epiphytes, endophytes or latent pathogens. UHPLC–Q-TOF-MS proved to be very effective as an analytical method for its rapidity, sensitivity and robustness.

This study provided new insights into several theoretical and practical aspects concerning the biology and epidemiology of *A. alternata, C. gloeosporioides* and *P. digitatum* and the identification of secondary metabolites these fungi produce during the interaction with the host plant. In particular, this is the first study addressing the effects of hailstorms on fruit contamination by mycotoxins produced by fungi residing in the fruit peel as latent pathogens. In the light of the results of this study, the risk of contamination appears low for metabolites of *A. alternata* and *C. gloeosporioides*, even on fruits injured by the hail. Conversely, it seems real in fruits infected by *P. digitatum* as in both the peel and juice of decayed fruits we found considerable amounts of *Penicillium*-related toxins such as Patulin and Rubratoxin B. Although mummified or severely blemished fruits usually do not enter the citrus fresh fruit supply chain as they are discarded before, this does not completely exclude the risk of contamination due to post-harvest rots. Moreover, mummified or severely blemished fruits are often present in bulks destined to the juice industry. An accurate and severe selection of these fruits would prevent or reduce the risk of contamination by micotoxins in the citrus juice industry.

## Author Contributions

Conceptualization, S.O.C., E.I.R., M.R.; methodology, S.O.C., M.R., E.I.R., C.L., G.M.; software, C.L., M.R.,E.I.R. and F.L.S.; validation, S.O.C., G.M., M.R., E.I.R, C.L. and F.L.S; formal analysis, M.R., E.I.R. and C.L.; investigation, E.I.R., M.R., and F.L.S.; resources, S.O.C.; data curation, M.R. and E.I.R; writing—original draft preparation, E.I.R., M.R. and F.L.S.; writing—review and editing, S.O.C., M.R..; visualization, S.O.C.; supervision, S.O.C and M.R. All authors have read and agreed to the published version of the manuscript.

## Funding

This research was funded by the University of Catania, Italy “Investigation of phytopathological problems of the main Sicilian productive contexts and eco-sustainable defense strategies (ME-DIT-ECO)” PiaCeRi-PIAno di inCEntivi per la Ricerca di Ateneo 2020-22 linea 2” “5A722192155”, by the projects “Smart and innovative packaging, postharvest rot management, and shipping of organic citrus fruit (BiOrangePack)” under the Partnership for Research and Innovation in the Mediterranean Area (PRIMA)—H2020 (E69C20000130001), the “Italie–Tunisie Cooperation Program 2014–2020” project “PROMETEO «Un village transfrontalier pour protéger les cultures arboricoles méditerranéennes en partageant les connaissances» cod. C-5-2.1-36, CUP 453E25F2100118000 and by European Union (NextGeneration EU), through the MUR-PNRR project SAMOTHRACE (ECS00000022).

## Data Availability Statement

Data will be available upon specific request to the authors.

## Acknowledgments

The authors wish also to thank Anna Davies for the English revision of the text

## Conflicts of Interest

The authors declare no conflict of interest.

## Disclaimer/Publisher’s Note

The statements, opinions and data contained in all publications are solely those of the individual author(s) and contributor(s) and not of MDPI and/or the editor(s). MDPI and/or the editor(s) disclaim responsibility for any injury to people or property resulting from any ideas, methods, instructions or products referred to in the content.

## Notes

### Competing Interest Statement

The authors have declared no competing interest.

## References

1. Ismail, M.; Zhang, J. (2004). Post-harvest citrus diseases and their control. Outlooks Pest Manag. 2004, *15*, 29.

2. Wu, G.A.; Terol, J.; Ibanez, V.; López-García, A.; Pérez-Román, E.; Borredá, C.; Domingo, D.; Tadeo, F.R.; Carbonell-Caballero, J.; Alonso, R.;, et al. Genomics of the origin and evolution of citrus. Nature 2018, 554, 311–316.

3. Aslam, K.; Iqbal, S.Z.; Razis, A.F.A.; Usman, S.,; Ali N.B. Patulin contamination of citrus fruits from Punjab and Northern Pakistan and estimation of associated dietary intake. Int. J. Env. Res. Pub. He. 2021, 18, 2270.

4. FAO (2020) Citrus fruit statistical compendium Rome 2021. Available online: https://www.fao.org/publications/card/fr/c/CB6492EN/ (Accessed on 8 June 2022).

5. Li, L.; Lyall, G.K.; Martinez-Blazquez, J.A.; Vallejo, F.A.; Tomas-Barberan, F.; Birch, K.M.; Boesch, C. Blood orange iuice consumption increases flow-mediated dilation in adults with overweight and obesity: a randomized controlled trial. J. Nutr. 2020, 150, 2287–2294.

6. Abobatta, W.F. Nutritional benefits of citrus fruits. Am. J. Biomed. Sci. Res. 2019, 3, 303–306.

7. Kahramanoğlu, İ.; Nisar, M.F.; Chen, C.; Usanmaz, S.; Chen, J.; Wan, C.. Light: An alternative method for physical control of postharvest rotting caused by fungi of citrus fruit. J. Food Qual. 2020, 1–12. Article ID 8821346.

8. Grosso, G.; Galvano, F.; Mistretta, A.; Marventano, S.; Nolfo, F.; Calabrese, G.; Buscemi, S.; Drago, F.; Veronesi, U.; Scuderi, A.. Red orange: experimental models and epidemiological evidence of its benefits on human health. Oxid. Med. Cell Longev. 2013, 2013, 157240.

9. Giménez-Sanchis, A.; Zhong, K.; Pintor, A.; Farina, V.; Besada, C. Understanding blood versus blond orange consumption: a cross-cultural study in four countries. Foods 2022, 11, 2686.

10. Akimitsu, K.; Peever, T.L.; Timmer, L.W. Molecular, ecological and evolutionary approaches to understanding Alternaria diseases of citrus. Mol. Plant Pathol. 2003, 4, 435–446.

11. Ramos, A.P.; Talhinhas, P.; Sreenivasaprasad, S.; Oliveira, H. Characterization of *Colletotrichum gloeosporioides*, as the main causal agent of citrus anthracnose, and *C. karstii* as species preferentially associated with lemon twig dieback in Portugal. Phytoparasitica 2016, 44, 549–561.

12. El boumlasy, S.; La Spada, F.; Tuccitto, N.; Marletta, G.; Mínguez, C.L.; Meca, G.; Rovetto, E.I.; Pane, A.; Debdoubi, A.; Cacciola, S.O. Inhibitory activity of shrimp waste extracts on fungal and oomycete plant pathogens. Plants 2021, 10, 2452.

13. Strano, M.C., Altieri, G.; Allegra, M.; Di Renzo, G.C.; Paterna, G.; Matera, A.; Genovese, F. Post-harvest technologies of fresh citrus fruit: advances and recent developments for the loss reduction during handling and storage. Horticulturae 2022, 8, 612.

14. Palou, L.; Usall, J.; Smilanick, J.L.; Aguilar, M.-J., Viñas, I. Evaluation of food additives and low-toxicity compounds as alternative chemicals for the control of *Penicillium digitatum* and *Penicillium italicum* on citrus fruit. Pest Manag. Sci. 2002, 58, 459–466.

15. Qi, J.; Pang, Y.; An, P.; Jiang, G.; Kong, Q.; Ren, X. Determination of metabolites of *Geotrichum citri-aurantii* treated with peppermint oil using liquid chromatography-mass spectrometry and gas chromatography-mass spectrometry. J. Food Biochem. 2019, 43, e12745.

16. Huang, F.; Chen, G.Q.; Hou, X.; Fu, Y.S.; Cai, L.; Hyde, K.D.; Li, H.Y. *Colletotrichum* species associated with cultivated citrus in China. Fungal Divers. 2013, 61, 61–74.

17. Riolo, M.; Aloi, F.; Pane, A.; Cara, M.; Cacciola S.O. Twig and shoot dieback of citrus, a new disease caused by *Colletotrichum* species. Cells 2021, 10, 449.

18. Wang, W.; de Silva, D.D.; Moslemi, A.; Edwards, J.; Ades, P.K.; Crous, P.W.; Taylor, P.W.J. Colletotrichum species causing anthracnose of citrus in Australia. J. Fungi 2021, 7,47.

19. Lopes da Silva, L.; Alvarado Moreno, H.L; Nunes Correia, H.L.; Ferreira Santana, M.; Vieira de Queiroz, M. Colletotrichum: species complexes, lifestyle, and peculiarities of some sources of genetic variability. Appl. Microbiol. Biot. 2020, 104, 1891–1904.

20. Woudenberg, J.H.C.; Seidl, M.F.; Groenewald, J.Z.; de Vries, M.; Stielow, J.B.; Thomma, B.P.H.J.; Crous, P.W. *Alternaria* section *Alternaria*: species, formae speciales or pathotypes? Stud. Mycol. 2015, 82, 1–21.

21. Lawrence, D.P.; Rotondo, F.; Gannibal, P.B. Biodiversity and taxonomy of the pleomorphic genus *Alternaria*. Mycol. Prog. 2016, 15. 1–22.

22. Patriarca, A. *Alternaria* in food products. Curr. Opin. Food Sci. 2016, 11, 1–9.

23. Timmer, L.V.; Peever, T.L.; Solel, Z.; Akimitzu, K. Alternaria diseases of citrus – Novel pathosystems. Phytopathol. Medit. 2003, 42, 99–112.

24. Garganese, F.; Schena, L.; Siciliano, I.; Prigigallo, M.I., Spadaro, D.; De Grassi, A.; Ippolito, A.; Sanzani, S.M. Characterization of citrus-associated *Alternaria* species in Mediterranean Aareas. PLoS One 2016, 16, e0163255.

25. Garganese, F.; Ippolito, A.; di Rienzo, V.; Lotti, C.; Montemurro, C.; Sanzani S.M. A new high-resolution melting assay for genotyping *Alternaria* species causing citrus brown spot. J. Sci. Food Agric. 2018, 98, 4578–4583.

26. Peever, T.L.; Su, G.; Carpenter-Boggs, L.; Timmer L.W. Molecular systematics of citrus-associated *Alternaria* species. Mycologia 2004, 96, 119–134.

27. Peever, T.L.; Carpenter-Boggs, L.; Timmer, L.W.; Carris, L.M.; Bhatia A. Citrus black rot is caused by phylogenetically distinct lineages of *Alternaria alternata*. Phytopathology 2005, 95, 512–518.

28. Tsitsigiannis, D.I.; Dimakopoulou, M.; Antoniou, P.P.; Tjamos, E.C. Biological control strategies of mycotoxigenic fungi and associated mycotoxins in Mediterranean basin crops. Phytopathol. Medit. 2012, 51, 158–174.

29. Bartholomew, H.P.; Bradshaw, M.; Jurick, W.M.; Fonseca, J.M. The good, the bad, and the ugly: Mycotoxin production during postharvest decay and their influence on tritrophic host–pathogen–microbe interactions. Front. Microbiol. 2021, 12, 611881.

30. Leslie, J.F.; Moretti, A.; Mesterházy, Á.; Ameye, M.; Audenaert, K.; Singh, P.K.; Richard-Forget, F.; Chulze, S.N.; Ponte, E.M.D.; Chala, A.;, et al. Key global actions for mycotoxin management in wheat and other small grains. Toxins 2021, 13, 725.

31. Nan, M.; Xue, H.; Bi, Y. Contamination, detection and control of mycotoxins in fruits and vegetables. Toxins 2022, 14, 309.

32. Wang, B.; Nong, X.H.; Zeng, W.N.; Li, S.S.; Li, G.Y.; Liu, J., … & Zheng, C. J. Study on bioactive secondary metabolites from the mangrove-derived fungus *Penicillium verruculosum* TGM14. Chem. Nat. Compd. 2022, 58, 812–815.

33. Schiff Jr., P.L. Ergot and its alkaloids. Am. J. Pharm. Educ. 2006, 70, 98.

34. Aloi F.; Riolo, M.; Sanzani, S.M.; Mincuzzi, A.;, Ippolito, A.; Siciliano, I.; Pane, A.; Gullino, M.L.; Cacciola, S.O. Characterization of *Alternaria* species associated with heart rot of pomegranate fruit. J. Fungi 2021, 7, 172.

35. Bentivenga, G.; Spina, A.; Ammar, K.; Allegra, M.; Cacciola, S.O. Screening of durum wheat (*Triticum turgidum* L. subsp. *durum* (Desf.) Husn.) Italian cultivars for susceptibility to Fusarium Head Blight incited by Fusarium graminearum. Plants 2021, 10, 68.

36. Stracquadanio, C.; Luz, C.; La Spada, F.; Meca, G.; Cacciola, S. O. Inhibition of mycotoxigenic fungi in different vegetable matrices by extracts of Trichoderma species. J. Fungi 2021, 7, 445.

37. Magan, N.; Olsen, M. (Eds.). Mycotoxins in food: detection and control; Woodhead Publishing Ltd, Abington (Cambridge), UK, 2004; p. 471.

38. Fernández-Cruz, M.L.; Mansilla, M.L.; Tadeo, J.L. (2010). Mycotoxins in fruits and their processed products: Analysis, occurrence and health implications. J. Adv. Res. 2010, *1*, 113–122.

39. Barkai-Golan, R.; Paster, N. (Eds.). Mycotoxins in fruits and vegetables, 1st ed.; Academic Press: san Diego, CA, USA, 2008; p.408.

40. Pallarés, N.; Carballo, D.; Ferrer, E.; Fernández-Franzón, M.; Berrada, H. Mycotoxin dietary exposure assessment through fruit juices consumption in children and adult population. Toxins 2019, 11, 684.

41. Carballo, D.; Pinheiro-Fernandes-Vieira, P.; Font, G.; Berrada, H.; Ferrer, E. Dietary exposure to mycotoxins through fruits juice consumption. Rev. de Toxicol. 2018, 35, 2–6

42. Spadaro, D.; Garibaldi, A.; Gullino, M.L. Occurrence of patulin and its dietary intake through pear, peach, and apricot juices in Italy. Food Addit. Contam. 2008, 1, 134–139.

43. Marino, A.; Nostro, A.; Fiorentino, C. (2009). Ochratoxin A production by *Aspergillus westerdijkiae* in orange fruit and juice. Int. J. Food Microbiol. 2009, *132*, 185–189.

44. Zouaoui, N.; Sbaii, N.; Bacha, H.; Abid-Essefi, S. Occurrence of patulin in various fruit juice marketed in Tunisia. Food Control 2015, 51, 356–360.

45. Juan, C.; Mañes, J.; Font, G.; Juan-García, A. (2017). Determination of mycotoxins in fruit berry by-products using QuEChERS extraction method. LWT - Food Science and Technology 2017, 86, 344–351.

46. Li, X.; Li, H.; Ma, W.; Guo, Z.; Li, X.; Li, X.; Zhang, Q. Determination of patulin in apple juice by single-drop liquid-liquid-liquid microextraction coupled with liquid chromatography-mass spectrometry. Food Chem. 2018, 257, 1–6

47. Groopman, J.D.; Kensler, T.W. Food safety: Mycotoxins. In Encyclopedia of Human Nutrition, 2nd ed.; Caballero, B., Allen, L., Prentice, A., Eds.; Academic Press, Cambridge, MA, USA, 2005; pp. 317–323.

48. Groopman, J.D.; Kensler, T.W.; Wild, C.P. Carcinogenesis in developing countries. Annu. Rev. Publ. Health 2008, 29, 187–203.

49. Wu, F.; Groopman, J.D.; Pestka, J.J. Public health impacts of foodborne mycotoxins. Annu. Rev. Food Sci. T. 2014, 5, 351–372.

50. EUR-Lex. Commission Regulation (EC) No 669/2009 of 24 July 2009 implementing Regulation (EC) No 882/2004 of the European Parliament and of the Council as regards the increased level of official controls on imports of certain feed and food of non-animal origin and amending Decision 2006/504/EC (Text with EEA relevance). Available online: http://data.europa.eu/eli/reg/2009/669/oj (Accessed on 7 May 2023).

51. Van de Perre, E.; Jacxsens, L.; Lachat, C.; El Tahan, F.; De Meulenaer, B. Impact of maximum levels in European legislation on exposure of mycotoxins in dried products: case of aflatoxin B1 and ochratoxin A in nuts and dried fruits. Food Chem. Toxicol. 2015, 75, 112–117.

52. Wang, M.; Jiang, N.; Xian, H.; Wei, D.; Shi, L.; Feng, X. A single-step solid phase extraction for the simultaneous determination of 8 mycotoxins in fruits by ultra-high performance liquid chromatography tandem mass spectrometry. J. Chromat. A 2016, 1429, 22–29.

53. EUR-Lex. Commission Regulation (EC) No 1881/2006 of 19 December 2006 setting maximum levels for certain contaminants in foodstuffs (Text with EEA relevance). Available online: http://data.europa.eu/eli/reg/2006/1881/oj (Accessed on 7 May 2023).

54. Hussain, S.; Asi, M.R.; Iqbal, M.; Khalid, N.; Wajih-ul-Hassan, S.; Ariño, A. Patulin mycotoxin in mango and orange fruits, juices, pulps, and jams marketed in Pakistan. Toxins 2020, 12, 52.

55. EUR-Lex. Commission Regulation (EU) No 165/2010 of 26 February 2010 amending Regulation (EC) No 1881/2006 setting maximum levels for certain contaminants in foodstuffs as regards aflatoxins (Text with EEA relevance). Available online: http://data.europa.eu/eli/reg/2010/165/oj (Accessed on 7 May 2023).

56. Boudergue, C.; Burel, C.; Dragacci, S.; Favrot, M. C.; Fremy, J.M.; Massimi, C.; Prigent, P.; Debongnie, P.; Pussemier, L.; Boudra, H.;, et al. Review of mycotoxin-detoxifying agents used as feed additives: mode of action, efficacy and feed/food safety. EFSA Supporting Publications 2009, 6, 22E.

57. EFSA Panel on Contaminants in the Food Chain (CONTAM). Schrenk, D.; Bodin, L.; Chipman, J.K.; del Mazo, J.; Grasl-Kraupp, B.; Hogstrand, C.; Hoogenboom, L.; Leblanc, J.-C.; Nielsen, E.; Ntzani, E.;, et al. Risk assessment of ochratoxin A in food. EFSA journal 2020, 18, e06113.

58. Escrivá, L.; Oueslati, S.; Font, G.; Manyes, L. *Alternaria* mycotoxins in food and feed: an overview. J. Food Quality 2017, Article ID 1569748.

59. Vaclavikova, M.; Dzuman, Z.; Lacina, O.; Fenclova, M.; Veprikova, Z.; Zachariasova, M.; Hajslova, J. Monitoring survey of patulin in a variety of fruit-based products using a sensitive UHPLC–MS/MS analytical procedure. Food Control 2015, 47, 577–584.

60. Murillo-Arbizu, M.; Amézqueta, S.; González-Peñas, E.; de Cerain, A.L. Occurrence of patulin and its dietary intake through apple juice consumption by the Spanish population. Food Chem. 2009, 113, 420–423.

61. Ruan, C.; Diao, X.; Zhang, H.; Zhang, L.; Liu, C. Development of a dispersive liquid–liquid microextraction technique for the analysis of citrinin, alternariol and alternariol monomethyl ether in fruit juices. Analytical Methods 2016, 8, 7944–7950.

62. Víctor-Ortega, M.D.; Lara, F.J.; García-Campaña, A.M.; del Olmo-Iruela, M. Evaluation of dispersive liquid–liquid microextraction for the determination of patulin in apple juices using micellar electrokinetic capillary chromatography. Food Control 2013, 31, 353–358

63. Zgoła-Grześkowiak, A.; Grześkowiak, T. Dispersive liquid-liquid microextraction. TrAC Trends Anal. Chem. 2011, 30, 1382–1399

64. Pallares, N.; Font, G.; Manes, J.; Ferrer, E. (2017). Multimycotoxin LC–MS/MS analysis in tea beverages after dispersive liquid–liquid microextraction (DLLME). J. Agric. Food Chem. 2017, *65*, 10282–10289.

65. Antep, H. M.; Merdivan, M. Development of new dispersive liquid−liquid microextraction technique for the identification of zearalenone in beer. Anal. Methods 2012, 4, 4129−4134.

66. Tolosa, J.; Font, G.; Mañes, J.; Ferrer, E. Multimycotoxin analysis in water and fish plasma by liquid chromatography-tandem mass spectrometry. Chemosphere 2016, 145, 402−408.

67. Rodríguez-Carrasco, Y.; Mañes, J.; Berrada, H.; Juan, C. Development and validation of a LC-ESI-MS/MS method for the determination of *Alternaria* toxins alternariol, alternariol methylether and tentoxin in tomato and tomato-based products. Toxins 2016, 8, 328.

68. Serrano, A.B.; Font, G.; Mañes, J.; Ferrer, E. Development a mitigation strategy of enniatins in pasta under home-cooking conditions. LWT-Food Sci. Technol. 2016, 65, 1017−1024

69. Gilbert, J.; Anklam, E. Validation of analytical methods for determining mycotoxins in food-stuffs. TrAC, Trends Anal. Chem. 2002, 21, 468–486.

70. Shi, H.; Li, S.; Bai, Y.; Prates, L.L.; Lei, Y.; Yu, P. Mycotoxin contamination of food and feed in China: occurrence, detection techniques, toxicological effects and advances in mitigation technologies. Food Control 2018, 91, 202–215.

71. Alshannaq, A.; Yu, J. H. Occurrence, toxicity, and analysis of major mycotoxins in food. Int. J. Env. Res. Pub. He. 2017, 14, 632.

72. Biselli, S.; Hummert C. Development of a multicomponent method for *Fusarium* toxins using LC–MS/MS and its application during a survey for the content of T-2 toxin and deoxynivalenol in various feed and food samples. Food Addit. Contam. 2005, 22, 752– 760.

73. Krska, R.; Welzig, E.; Boudra, H. Analysis of *Fusarium* toxins in feed. Anim. Feed Sci. Technol. 2007, 137, 241–264

74. Martins, H.M.; Mendes Guerra, M.M.; d’Almeida Bernardo F.M.. Occurrence of aflatoxin B1 in dairy cow feed over 10 years in Portugal (1995–2004). Rev. Iberoam. Micol. 2007, 24, 69–71.

75. Vrabcheva, T.; Usleber, E.; Dietrich, R.; Martlbauer, E. Co-occurrence of ochratoxin A and citrinin in cereals from Bulgarian villages with a history of Balkan endemic nephropathy. J. Agric. Food Chem. 2000, 48, 2483–2488.

76. Delmulle, B.; De Saeger, S.; Adams, A.; De Kimpe, N.; Van Peteghem, C. Development of a liquid chromatography/tandem mass spectrometry method for the simultaneous determination of 16 mycotoxins on cellulose filters and in fungal cultures. Rapid Commun. Mass Spectrom. 2006, 20, 771–776.

77. Kokkonen, M.; Jestoi, M.; Rizzo A. Determination of selected mycotoxins in mould cheeses with liquid chromatography coupled to tandem with mass spectrometry. Food Addit. Contam. 2005, 22, 449–456.

78. Sorensen, L.K.; Elbaek, T.H. Determination of mycotoxins in bovine milk by liquid chromatography tandem mass spectrometry. J. Chromatogr. B Analyt. Technol. Biomed. Life Sci. 2005, 820, 183–196.

79. Garcia-Villanova, R.J.; Cordon, C.; Gonzalez Paramas, A.M.; Aparicio, P.; Garcia Rosales M.E.. Simultaneous immunoaffinity column clean-up and HPLC analysis of aflatoxins and ochratoxin A in Spanish bee pollen. J. Agric. Food. Chem. 2004, 52, 7235–7239

80. Aresta A.; Cioffi, N.; Palmisano, F.; Zambonin, C.G. Simultaneous determination of ochratoxin A and cyclopiazonic, mycophenolic, and tenuazonic acids in cornflakes by solidphase micro-extraction coupled to high-performance liquid chromatography. J. Agric. Food Chem. 2003, 51, 5232–5237.

81. Chan, D.; MacDonald, S.J.; Boughtflower, V.; Brereton, P.. Simultaneous determination of aflatoxins and ochratoxin A in food using a fully automated immunoaffinity column cleanup and liquid chromatography-fluorescence detection. J. Chromatogr. A 2004,1059, 13–16

82. Rahmani, A.; Jinap, S.; Soleimany, F. Qualitative and quantitative analysis of mycotoxins. Compr. Rev. Food Sci. Food Saf. 2009, 8, 202–251.

83. Soleas, G.J.; Yan, J.; Goldberg, D.M. Assay of ochratoxin A in wine and beer by highpressure liquid chromatography photodiode array and gas chromatography mass selective detection. J. Agric. Food Chem. 2001, 49, 2733–2740.

84. He, Y.; Li, Z.; Wang, W.; Sooranna, S.R.; Shi, Y.; Chen, Y.; Wu, C.; Zeng, J.; Tang, Q.; Xie, H. Chemical Profiles and Simultaneous Quantification of *Aurantii fructus* by Use of HPLC-Q-TOF-MS Combined with GC-MS and HPLC Methods. Molecules 2018, 23, 2189.

85. Dopazo, V.; Luz, C.; Mañes, J.; Quiles, J.M.; Carbonell, R.; Calpe, J.; Meca, G. Bio-preservative potential of microorganisms isolated from red grape against food contaminant fungi. Toxins 2021, 13, 412.

86. Thurman, E.M.; Ferrer, I.; Fernández-Alba, A.R. Matching unknown empirical formulas to chemical structure using LC/MS TOF accurate mass and database searching: example of unknown pesticides on tomato skins. J. Chromat. A 2005, 1067, 127–134.

87. Xing, T.T.; Zhao, X.J.; Zhang, Y.D.; Li, Y.F. Fast separation and sensitive quantitation of polymethoxylated flavonoids in the peels of citrus using UPLC-Q-TOF-MS. J. Agric. Food Chem. 2017, 65, 2615–2627.

88. Wang, H.; Guo, Y.; Luo, Z.; Gao, L.; Li, R.; Zhang, Y.; Kalaji, H.M.; Qiang, S.; Chen, S. Recent advances in *Alternaria* phytotoxins: a review of their occurrence, structure, bioactivity, and biosynthesis. J. Fungi 2022, 8, 168.

89. Nakatsuka, S.I.; Ueda, K.; Goto, T.; Yamamoto, M.; Nishimura, S.; Kohmoto, K. Structure of AF-toxin II, one of the host-specific toxins produced by *Alternaria alternata* strawberry pathotype. Tetrahedron Lett. 1986, 27, 2753–2756.

90. Bensassi, F.; Gallerne, C.; Hajlaoui, M.R.; Bacha, H.; Lemaire, C. (2011). Mechanism of Alternariol monomethyl ether-induced mitochondrial apoptosis in human colon carcinoma cells. Toxicology 2011, *290*, 230–240.

91. Berestetskiy, A.O.; Dalinova, A.A.; Volosatova, N.S. Metabolite profiles and biological activity of extracts from *Alternaria sonchi* S-102 culture grown by different fermentation methods. Applied Biochem. Microbiol. 2019, 55, 284–293.

92. Debbab, A.; Aly, A.H.; Edrada-Ebel, R.; Wray, V.; Müller, W.E.; Totzke, F.; Zirrgiebel, U.; Schäctele, C.; Kubbutatt, M.H.G.;, et al. Bioactive metabolites from the endophytic fungus *Stemphylium globuliferum* isolated from *Mentha pulegium*. J. Nat. Prod. 2009, 72, 626–631.

93. Shaaban, M.; Shaaban, K.A.; Abdel-Aziz, M. S. Seven naphtho-γ-pyrones from the marine-derived fungus *Alternaria alternata*: structure elucidation and biological properties. Org. Med. Chem. Lett. 2012, 2, 1–8.

94. Meena, M.; Samal, S. (2019). *Alternaria* host-specific (HSTs) toxins: an overview of chemical characterization, target sites, regulation and their toxic effects. Toxicol. Rep. 2019, 6, 745–758.

95. Liu, Y.; Palaniveloo, K.; Alias, S.A.; Sathiya Seelan, J.S. Species diversity and secondary metabolites of *Sarcophyton*-associated marine fungi. Molecules 2021, 26, 3227.

96. Yang, Y.-S.; Johnson, D.R.; Dowler, W.M. Pathogenicity of *Alternaria angustiovoidea* on leafy spurge. Plant Dis. 1990, 74, 601–604.

97. Elkhateeb, W.A.; Kolaibe, A.G.A.; Elkhateeb, A.; Daba, G.M. Allergen, pathogen, or biotechnological tool? The dematiaceous fungi Alternaria what’s for it and what’s on it. J. Pharm. Pharmacol. Res. 2021, 4, 1–6.

98. Suemitsu, R.; Ohnishi, K.; Morikawa, Y.; Ideguchi, I.; Uno, H. (). Porritoxinol, a phytotoxin of *Alternaria porri*. Phytochemistry 1994, 35, 603–605.

99. Jiménez-Teja, D.; Daoubi, M.; Collado, I.G.; Hernández-Galán, R.. Lipase-catalyzed resolution of 5-acetoxy-1, 2-dihydroxy-1, 2, 3, 4-tetrahydronaphthalene. Application to the synthesis of (+)-(3R, 4S)-cis-4-hydroxy-6-deoxyscytalone, a metabolite isolated from Colletotrichum acutatum. Tetrahedron 2009, 65, 3392-3396.

100. Yang, Z.D.; Li, Z.J.; Zhao, J.W.; Sun, J.H.; Yang, L.J.; Shu, Z.M. Secondary metabolites and PI3K inhibitory activity of Colletotrichum gloeosporioides, a fungal endophyte of *Uncaria rhynchophylla*. Curr. Microbiol. 2019, 76, 904–908.

101. Chakraborty, A.; Ray, P. Mycoherbicides for the noxious meddlesome: can *Colletotrichum* be a budding candidate?. Front. Microbiol. 2021, 12, 754048.

102. Moraga, J.; Gomes, W.; Pinedo, C., Cantoral, J.M.; Hanson, J.R.; Carbú, M.; Garrido, C.; Duran-Patron, R.; Collado, I.G. The current status on secondary metabolites produced by plant pathogenic *Colletotrichum* species. Phytochem. Rev. 2019, 18, 215–239.

103. Kim, J. W., Shim, S. H. (2019). The fungus *Colletotrichum* as a source for bioactive secondary metabolites. Arch. Pharm. Res. 2019, *42*, 735–753.

104. Gohbara, M.; Kosuge, Y.; Yamasaki, S.; Kimura, Y.; Suzuki, A.; Tamura, S. Isolation, structures and biological activities of colletotrichins, phytotoxic substances from *Colletotrichum nicotianae*. Agr. Biol. Chem. Tokyo 1978, 42, 1037–1043.

105. Inoue, M.; Takenaka, H.; Tsurushima, T.; Miyagawa, H.; Ueno, T. Colletofragarones A1 and A2, novel germination self-inhibitors from the fungus *Colletotrichum fragariae*. Tetrahedron Lett. 1996, 37, 5731–5734.

106. Aoyagi, A.; Ito-Kobayashi, M.; Ono, Y.; Furukawa, Y.; Takahashi, M.; Muramatsu, Y.; Umetani, M.; Takatsu, T. Colletoic acid, a novel 11β-hydroxysteroid dehydrogenase type 1 inhibitor from *Colletotrichum gloeosporioides* SANK 21404. J. Antibiot. 2008, 61, 136–141.

107. Hsiao, Y.; Cheng, M.J.; Chang, H.S.; Wu, M.D.; Hsieh, S.Y.; Liu, T.W.; Lin, C.H.; Yuan, G.F.; Chen I.S. Six new metabolites produced by *Colletotrichum aotearoa* 09F0161, an endophytic fungus isolated from *Bredia oldhamii*. Nat. Prod. Res. 2016, 30, 251–258.

108. H.; Root, N.; Jabeen, F.; Al-Harrasi, A.; Al-Rawahi, A.; Ahmad, M.; Hassan, Z.; Abbas, G.; Mabood, F.; Shah, A.;, et al. Seimatoric acid and colletonoic acid: two new compounds from the endophytic fungi, Seimatosporium sp. and Colletotrichum sp. Chinese Chem. Lett. 2014, 25, 1577–1579.

109. Wei, B.; Yang, Z.D.; Chen, X.W.; Zhou, S.Y.; Yu, H.T.; Sun, J.Y.; Yao, X.J.; Wang, Y.G.; Xue, H.Y. Colletotrilactam A–D, novel lactams from *Colletotrichum gloeosporioides* GT-7, a fungal endophyte of *Uncaria rhynchophylla*. Fitoterapia 2016, 113, 158–163.

110. Tianpanich, K.; Prachya, S.; Wiyakrutta, S.; Mahidol, C.; Ruchirawat, S.; Kittakoop, P. Radical scavenging and antioxidant activities of isocoumarins and a phthalide from the endophytic fungus *Colletotrichum* sp. J. Nat. Prod. 2011, 74, 79–81.

111. Yang, Z.; Bao, L.; Yin, Y.; Ding, G.; Ge, M.; Chen, D.; Qian, X. Pyrenocines N–O: two novel pyrones from Colletotrichum sp. HCCB03289. J. Antibiot. 2014, 67, 791-793.

112. Ichihara, A.; Murakami, K.; Sakamura, S.. Synthesis of pyrenocines A, B and pyrenochaetic acid A. Tetrahedron 1987, 43, 5245–5250.

113. Nunez, F.; Westphal, C.D.; Bermudez, E.; Asensio, M.A. Production of secondary metabolites by some terverticillate penicillia on carbohydrate-rich and meat substrates. J. Food Prot. 2007, 70, 2829–2836.

114. Frisvad, J.C.; Smedsgaard, J.; Larsen, T.O.; Samson, R.A. Mycotoxins, drugs and other extrolites produced by species in *Penicillium* subgenus *Penicillium*. Stud. Mycol. 2004, 49, e41.

115. Ali, H.; Ries, M.I.; Lankhorst, P.P.; van der Hoeven, R.A.; Schouten, O.L.; Noga, M.; Hankemeier, T.; van Peij, N.N.M.E.; Bovenberg, R.A.L.; Vreeken, R.B.;, et al. A non-canonical NRPS is involved in the synthesis of fungisporin and related hydrophobic cyclic tetrapeptides in *Penicillium chrysogenum*. PLoS One 2014, 9, e98212.

116. Wang, L.; Zhou, H.B.; Frisvad, J.C.; Samson, R.A. *Penicillium persicinum*, a new griseofulvin, chrysogine and roquefortine C producing species from Qinghai province, China. Antonie van Leeuwenhoek 2004, 86, 173–179.

117. Frisvad, J.C.; Samson, R.A.; Rassing, B.R.; van der Horst, M.I.; Van Rijn, F.T.J.; Stark, J. *Penicillium discolor*, a new species from cheese, nuts and vegetables. Antonie van Leeuwenhoek 1997, 72, 119–126.

118. Das, T.; Ray, P.; Nandy, S.; Al-Tawaha, A.R.; Pandey, D.K.; Kumar, V.; Dey, A. Piezophilic fungi: sources of novel natural products with preclinical and clinical significance. In *Extremophilic Fungi: Ecology, Physiology and Applications*; Sahay, S., Ed.; Springer, Singapore, 2022; pp. 523–545.

119. Ortega, H.E.; Torres-Mendoza, D.; Caballero E, Z.; Cubilla-Rios, L. (2021). Structurally uncommon secondary metabolites derived from endophytic fungi. J. Fungi 2021, *7*, 570.

120. Zain, M. E. Effect of olive oil on secondary metabolite and fatty acid profiles of *Penicillium expansum*, *Aspergillus flavus*, A. parasiticus and A. ochraceus. Aust. J. Basic Appl. Sci. 2009, 3, 4274–4280.

121. Ariza, M.R.; Larsen, T.O.; Petersen, B.O.; Duus, J.Ø.; Christophersen, C.; Barrero, A.F. A novel alkaloid serantrypinone and the spiro azaphilone daldinin D from *Penicillium thymicola*. J. Nat. Prod. 2001, 64, 1590–1592.

122. Larsen, T.O.; Gareis, M.; Frisvad, J.C. Cell cytotoxicity and mycotoxin and secondary metabolite production by common penicillia on cheese agar. J. Agric. Food Chem. 2002, 50, 6148–6152.

123. Evidente, A.; Kornienko, A.; Cimmino, A.; Andolfi, A.; Lefranc, F.; Mathieu, V.; Kiss, R. Fungal metabolites with anticancer activity. Nat. Prod. Rep. 2014, 31, 617–627.

124. Perrone, G.; Susca, A. Penicillium species and their associated mycotoxins. In Mycotoxigenic Fungi; Springer: New York, NY, USA, 2017; pp. 107–119.

125. Riolo, M.; Pane, A.; Santilli, E.; Moricca, S.; Cacciola, S. O. Susceptibility of Italian olive cultivars to various Colletotrichum species associated with fruit anthracnose. Plant Pathology, 2023, 72(2), 255–267.

126. Awuchi, C.G.; Ondari, E.N.; Ogbonna, C.U.; Upadhyay, A.K.; Baran, K.; Okpala, C.O.R.; Korzeniowska, M.; Guiné, R.P.F. Mycotoxins affecting animals, foods, humans, and plants: types, occurrence, toxicities, action mechanisms, prevention, and detoxification strategies—A revisit. Foods 2021, 10, 1279.

127. Pallarés, N.; Carballo, D.; Ferrer, E.; Fernández-Franzón, M.; Berrada, H. Mycotoxin dietary exposure assessment through fruit juices consumption in children and adult population. Toxins 2019, 11, 684.

128. Müller M.E.; Steinhaus M.; Böhm J.; Müller A.; Langer E.; Andratsch M.; Mack B.; Novohradská E.; Klempier N. In vitro anti-inflammatory and anti-oxidative effects of altenariol and altertoxin II isolated from *Alternaria* species. Toxins 2018, 10, 228.

129. Mitchell, C.G.; Slight, J.; Donaldson, K. Diffusible component from the spore surface of the fungus *Aspergillus fumigatus* which inhibits the macrophage oxidative burst is distinct from gliotoxin and other hyphal toxins. Thorax 1997, 52, 796–801.

130. Tolosa, J.; Barba, F.J.; Pallarés, N.; Ferrer, E. Mycotoxin identification and in silico toxicity assessment prediction in Atlantic Salmon. Mar. Drugs 2020, 18, 629.

131. da Cruz Cabral, L.; Rodriguero, M.; Stenglein, S.; Nielsen, K. F.; Patriarca, A. Characterization of small-spored *Alternaria* from Argentinean crops through a polyphasic approach. Int. J. Food Microbiol. 2017, 257, 206–215.

132. Man, Y.; Liang, G.; Li, A.; Pan, L. Analytical methods for the determination of *Alternaria* mycotoxins. Chromatographia 2017, 80, 9–22

133. Mirocha, C.J.; Gilchrist, D.G.; Shier, W.T.; Abbas, H. K.; Wen, Y.; Vesonder R.F. AAL Toxins, funionisms (biology and chemistry) and host-specificity concepts. Mycopathologia, 1992, 117, 47–56.

134. Yamagishi, D.; Akamatsu, H.; Otani, H.; Kodama M. Pathological evaluation of host-specific AAL-toxins and fumonisin mycotoxins produced by *Alternaria* and *Fusarium* species. J. Gen. Plant Pathol. 2006, 72, 323–326.

135. Tsuge, T.; Harimoto, Y.; Akimitsu, K.; Ohtani, K.; Kodama, M.; Akagi, Y.; Egusa, M.; Yamamoto, M.; Otani, H. Host-selective toxins produced by the plant pathogenic fungus *Alternaria alternata*. FEMS Microbiol. Rev. 2013, 37, 44–66.

136. Xu, J.; Yang, X.; Lin, Q. Chemistry and biology of *Pestalotiopsis*-derived natural products. Fungal Divers. 2014, 66, 37–68.

137. Evidente, A.; Punzo, B.; Andolfi, A.; Berestetskiy, A.; Motta, A. Alternethanoxin A and B, polycyclic ethanones produced by *Alternaria sonchi*, potential mycoherbicides for *Sonchus arvenis* Biocontrol. J. Agric. Food Chem. 2009, 57, 6656−6660.

138. Riolo, M.; Luz, C.; Santilli, E.; Meca, G.; Cacciola, S. O. Secondary metabolites produced by Colletotrichum spp. on different olive cultivars. bioRxiv, 2022, 2022-11.

139. Sadahiro, Y.; Hitora, Y.; Kimura, I.; Hitora-Imamura, N.; Onodera, R.; Motoyama, K.; & Tsukamoto, S. Colletofragarone A2 Inhibits Cancer Cell Growth In Vivo and Leads to the Degradation and Aggregation of Mutant p53. Chemical Research in Toxicology, 2022, 35(9), 1598–1603.

140. Yan, X.; Qi, M.; Li, P.; Zhan, Y.; Shao, H. Apigenin in cancer therapy: Anti-cancer effects and mechanisms of action. Cell Biosci. 2017, 7, 50.

141. El Hajj Assaf, C.; Zetina-Serrano, C.; Tahtah, N.; Khoury, A.E.; Atoui, A.; Oswald, I.P.; Puel, O.; Lorber, S. Regulation of Secondary Metabolism in the *Penicillium* Genus. Int. J. Mol. Sci. 2020, 21, 9462

142. Hussain, S.; Asi, M.R.; Iqbal, M.; Khalid, N.; Wajih-ul-Hassan, S.; Ariño, A. Patulin Mycotoxin in Mango and Orange Fruits, Juices, Pulps, and Jams Marketed in Pakistan. Toxins 2020, 12, 52.

143. Moake, M.M.; Zakour, O.I.P.; Worobo, R.W. Comprehensive review of patulin control methods in food. Compr. Rev. Food Sci. Food Saf. 2005, 4, 8–21.

144. Wogan, G. N.; Edwards, G. S.; Newberne, P. M. Acute and chronic toxicity of rubratoxin B. Toxicology and applied pharmacology, 1971,19(4), 712-720.

145. Yabe, K.; Nakamura, S.; Nakajima, M.; Fujimoto, H. Isolation and characterization of rubratoxin B biosynthetic intermediates from Penicillium sp. TP-F0595. J. Antibiot. 2003, 56, 534-540.

146. Ding, Z.; Tao, T.; Wang, L.; Zhao, Y.; Huang, H.; Zhang, D.; Liu, M.; Wang, Z.; Han, J. Bio-prospecting of Novel and Bioactive Metabolites from Endophytic Fungi Isolated from Rubber Tree Ficus elastica Leaves. Microbiol. Biotechnol. 2019, 29, 731–738

147. Ren-Yi, G.; Lei, X.; Yi, K.; Iii-Ming, C.; Jian-Chun, Q.; Li, L.; Sheng-Xiang, Y.; Li-Chun, Z. Chaetominine,(+)-alantrypinone, questin, isorhodoptilometrin, and 4-hydroxybenzaldehyde produced by the endophytic fungus Aspergillus sp. YL-6 inhibit wheat (Triticum aestivum) and radish (Raphanus sativus) germination. J. Plant Interact. 2015, 10, 87-92.

148. Kuriyama, T.; Kakemoto, E.; Takahashi, N.; Imamura, K. I.; Oyama, K.; Suzuki, E.; Harimaya, K.; Yaguchi, T.; Ozoe, Y. Receptor assay-guided isolation of anti-GABAergic insecticidal alkaloids from a fungal culture. J. Agric. Food Chem. 2004, 52, 3884–3887.

149. Zhu, X.; Zhou, D.; Liang, F.; Wu, Z.; She, Z.; Li, C. enochalasin K, a new unusual chaetoglobosin from the mangrove endophytic fungus Penicillium chrysogenum V11 and its effective semi-synthesis. Fitoterapia 2017, 123, 23–28

150. Masi, M.; Aloi, F.; Nocera, P.; Cacciola, S.O.; Surico, G.; Evidente, A. Phytotoxic metabolites isolated from *Neufusicoccum batangarum*, the causal agent of the scabby canker of cactus pear (Opuntia ficus-indica L.). Toxins 2020, 12, 126.

151. Pryor, B.M.; Michailides, T.J. Morphological, pathogenic, and molecular characterization of *Alternaria* isolates associated with Alternaria late blight of pistachio. Phytopathology 2002, 92, 406–416.

152. Simmons, E.G. Alternaria*: an identification manual: fully illustrated and with catalogue raisonné 1796–2*007; CBS Fungal Biodiversity Centre: Utrecht, The Netherlands, 2007; p. 775.

153. Agrios, G. Plant Pathology. 5th ed.; Elsevier, Academic Press, Amsterdam, NL, 2005; pp. 26–27, 398-401.

154. White, T.J.; Bruns, T.; Lee, S.; Taylor, J.W. Amplification and direct sequencing of fungal ribosomal RNA genes for phylogenetics. In PCR protocols: a guide to methods and applications; Innis, M.A., Gelfand, D.H., Sninsky, J.J., White, T.J., Eds.; Academic Press, Inc.: San Diego, CA, USA, 1990; Volume 18, pp. 315–322.

155. Carbone, I.; Kohn, L.M. A method for designing primer sets for speciation studies in filamentous ascomycetes. Mycologia 1999, 91, 553–556.

156. Glass, N.L.; Donaldson, G.C. Development of primer sets designed for use with the PCR to amplify conserved genes from filamentous ascomycetes. Appl. Environ. Microbiol. 1995, 61, 1323–30.

157. Hong, S.G.; Cramer, R.A.; Lawrence, C.B.; Pryor, B.M. Alt a 1 allergen homologs from *Alternaria* and related taxa: analysis of phylogenetic content and secondary structure. Fungal Genet. Biol. 2005, 42, 119–129.

158. Widiantini, F.; Yulia, E.; Nasahi, C.. Antifungal activity of methanol-extracted secondary metabolites of rhizobacteria isolated from rhizosphere of oil palm trees against *Ganoderma boninense* Pat. IOP Conference Series: Earth Environ. Sci. 2019, 334, 012037.

159. El Jai, A.; Juan, C.; Juan-Garcia, A.; Manes, J.; Zinedine, A. Multi-mycotoxin contamination of green tea infusion and dietary exposure assessment in Moroccan population. Food Res. Intern. 2021, 140, 109958.

160. Pang, Z.; Chong, J.; Zhou, G.; De Lima Morais, D.A.; Chang, L.; Barrette, M.; Gauthier, C.; Jacques, P.É.; Li, S.; Xia, J. MetaboAnalyst 5.0: narrowing the gap between raw spectra and functional insights. Nucleic Acids Res. 2021, 49, W388–W396.

